# Multimodal Human Scalp Atlas Defines Cell Landscape and Lineage Architecture In Situ

**DOI:** 10.64898/2026.06.29.735239

**Authors:** Edward B. Li, Christopher M. Stephens, Meredith Klay, Adrian Carcamo, Jimin Han, Victor L. Quan, Maria L. Colavincenzo, Ross L. Pearlman, Simon S. Yoo, Dongmei Wang, Nicolas Tissot, Thomas Bornschlögl, Inês Sequeira, Rui Yi

## Abstract

Human scalp hair has an extraordinary ability to grow continuously for years while maintaining structural and functional integrity. However, the cell states and lineage organization that enable this capacity and how they are disrupted in inflammatory hair loss disorders remain poorly defined in humans. Here we establish a high-resolution, multimodal atlas of human scalp by integrating deep-coverage spatial transcriptomics with single-cell RNA-seq and multiomics data. This reference resolves spatially organized epithelial and mesenchymal states and links *in situ* transcriptional programs to chromatin accessibility dynamics and lineage trajectories at single-cell resolution, revealing human-specific principles of tissue organization and previously unrecognized features of hair follicle architecture and lineage progression. We validate key aspects of matrix cell organization and cell activities using live imaging, connecting molecularly defined cell states to dynamic cell behaviors and lineage progression in the matrix. Leveraging the atlas as a spatial reference, we project patient scRNA-seq profiles from alopecia areata and lichen planopilaris onto defined cell compartments, resolving disease-specific perturbations in fibroblasts, epithelial and immune populations. This comparison delineates distinct cellular programs associated with non-scarring versus scarring hair loss and highlights compartment- and state-specific pathways with diagnostic and therapeutic potential. Together, this work provides a foundational resource for human hair biology and establishes a generalizable framework for spatially resolved, multimodal interrogation of tissue organization and disease in complex human tissues.

## Main text

Knowledge of the cellular landscape and tissue architecture of human organs is essential for understanding how tissues form, maintain homeostasis, and repair themselves and how these processes go awry in disease. Advances in single-cell genomics, including single-cell RNA sequencing (scRNA-seq), single-cell multiomics, and spatial transcriptomics (ST), have transformed human tissue analysis by resolving cellular states and heterogeneity at unprecedented molecular depth for virtually all organs^1–6^. Yet, dissociation-based single cell sequencing approaches disrupt native tissue context, obscuring spatial and lineage relationships among closely related, transcriptionally similar populations. As a result, many morphologically recognizable, biochemically distinct and functionally important cell types have not been defined unambiguously. This incomplete view of *in situ* cellular organization hampers reconstruction of cell lineage relationships and interrogation of regulatory mechanisms that underpin important functions of human tissues in health and disease, including stem/progenitor self-renewal and differentiation. To address these challenges, both imaging- and sequencing-based ST techniques have been developed to enhance *in situ* analysis of cellular landscape^5–8^. However, technical constraints, such as a limited number of gene probes, low effective spatial resolution, and modest transcript capture efficiency per spot, have compromised robust single cell spatial mapping and confident inference of lineage organization in intact human tissues. Consequently, elucidating human cell lineages in their native microenvironment remains a major challenge.

In this study, we use the human scalp hair follicle (HF), an anatomically ordered mini-organ with well-defined spatial compartments, to map tissue architecture and lineage organization *in situ*. We establish an optimized experimental pipeline that generates high-resolution, whole-transcriptome ST maps of the human scalp. The deep transcript capture of our ST data enables stand-alone spatial lineage analysis and provides high-dimensional anchors for accurate co-embedding and integration with matched scRNA-seq and single-cell multiome data. The multimodal atlas delineates the cellular landscape of the scalp with spatial granularity and open chromatin context, resolving closely related keratinocyte and mesenchymal subpopulations that are difficult to distinguish by morphology alone. Leveraging these high-resolution data, we reconstruct the lineage architecture of the densely packed hair matrix, linking spatially organized rapid proliferation-to-differentiation waves to fate-poised chromatin accessibility at the root of each matrical lineage, and we validate key aspects of matrix organization and lineage progression using live imaging of intact human hair. Finally, by projecting scRNA-seq profiles from patients with alopecia areata and lichen planopilaris onto this reference atlas, we identify disease-associated alterations across distinct fibroblasts, epithelial and immune cell populations, revealing divergent cellular programs consistent with non-scarring and scarring inflammatory hair loss, respectively. Together, this work establishes a reference framework for human HF biology (https://www.hha.fsm.northwestern.edu) and provides a generalizable strategy for spatially resolved, multimodal analysis of complex human tissues in health and disease at single-cell resolution.

### High-resolution Atlas of Human Scalp Skin

Skin is the largest organ of the human body and displays remarkable regional specialization. The scalp is one of the most distinct human features, containing approximately 100,000 hair follicles (HFs). Each HF is a spatially organized mini-organ composed of diverse epithelial and mesenchymal cell populations embedded in complex, heterogenous microenvironments, and produces hair shaft at ∼0.3-0.4mm per day^9,10^ (Fig. 1a). To capture the cellular heterogeneity of healthy human scalp, we collected skin biopsies from healthy donors and freshly discarded scalp tissue from surgery in a sex-balanced, demographically diverse cohort (Fig. 1b and Supplementary Table 1). Specimens were visually inspected and follicles were confirmed to be in phenotypic anagen^11^ before single-cell dissociation and generation of scRNA-seq and single-nuclei multiome (snRNA+snATAC) libraries (Fig. 1b). After stringent quality control and dataset integration to correct batch effects while preserving biological heterogeneity (see Methods), we retained 128,185 high quality cells and nuclei for downstream analyses (Fig. 1c).

**Fig. 1:**
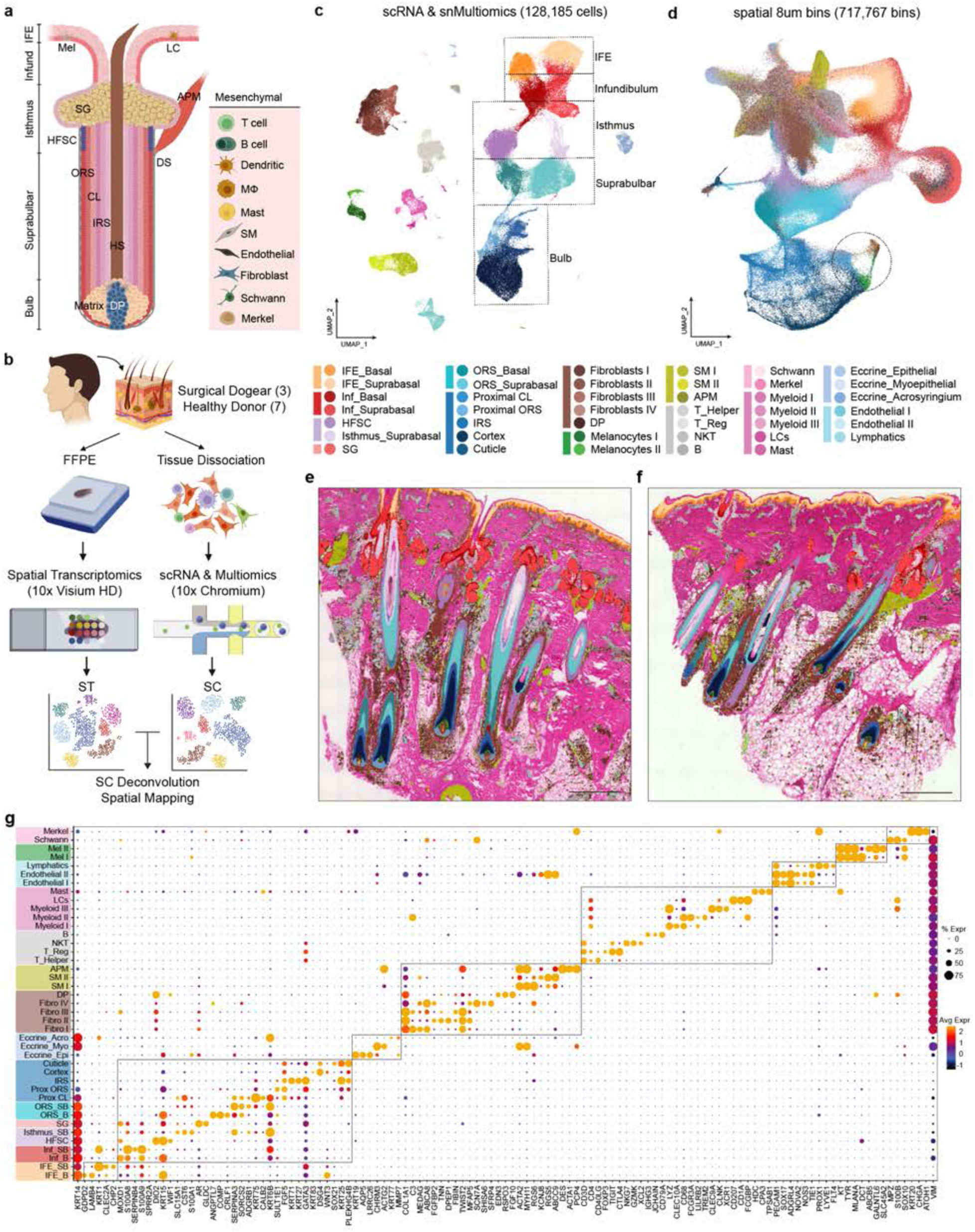
Single-Cell and Spatial Transcriptomic Atlas of the Human Scalp Skin. **(a)** Schematic of human anagen hair follicle and scalp skin cellular composition. **(b)** Experimental approach for sample collection and processing. **(c)** UMAP of the integrated single-cell RNA-seq and multiomics atlas of human scalp skin. Boxed: major keratinocyte populations extending from the interfollicular epidermis (IFE) through the hair follicle. **(d)** UMAP of 8µm spatial transcriptomic bins with cluster identities assigned based on SC label transfer, spatial marker genes, and histologic tissue localization cross-validation. Circle: dermal papilla-melanocytes-keratinocyte aggregation in the hair bulb/matrix. **(e-f)** Spatial plotting of atlas-level SC clusters on ST samples S2 (E) and S3 (F). Scale bar, 1mm. **(g)** Dot plot of key marker gene expression by atlas-level cluster identities.

Across the integrated dataset, cells mapped to 9 major types: keratinocytes, eccrine gland cells, fibroblasts, smooth muscle cells, lymphoid cells, myeloid cells, melanocytes, vascular cells, and neural cells (Extended Data Fig. 1a). High-resolution clustering resolved 41 sub-clusters spanning abundant populations and rare cell types, such as Merkel cells and dermal papilla (DP) cells. Even at this global level, numerous keratinocyte and stromal subtypes were readily distinguished, underscoring the remarkable cellular diversity of the scalp (Fig. 1c). Because we aimed to resolve lineage organization and tissue architectures within the hair bulb, which sustains anagen hair growth for up to a decade^12,13^, we further micro-dissected individual HFs below the isthmus and profiled bulb-containing portions in several matched samples (Extended Data Fig. 1b). Consistent with this physical separation, bulb preparations were devoid of cells from clusters representing the upper HF, interfollicular epidermis (IFE), sebaceous gland (SG), eccrine gland, and papillary (upper) dermis, while enriching for lower HF and DP populations (Extended Data Fig. 1b). These results indicate that our cell preparation, computational integration, and cell type annotation pipeline preserved the authentic biological partitioning and cellular heterogeneity of human scalp skin (Extended Data Fig. 1c).

While small collections of marker genes readily distinguish divergent cell types, many sub-populations of keratinocytes and mesenchymal cells occupy distinct spatial positions yet differ only subtly in transcriptional state and lack population exclusive markers. These subtle differences were also reflected by reduced cluster specificity in inter-cluster differential gene expression (DEG) analysis among closely related populations (Supplementary Table 2). Moreover, without spatial information, the native localization of these nuanced populations cannot be reliably defined. To provide spatial context and enable confident identification of fine-grained cell identities, we therefore performed sequencing-based spatial transcriptomics (ST) on matched scalp samples from healthy donors using the 10x Genomics Visium HD platform^14^.

A recent benchmarking study reported that the Visium HD platform, enabled by seamless tiling of genome-scale capture probes, outperforms Stereo-seq (220nm capture spot with a 500nm spot-to-spot pitch) in effective gene capture, transcript diffusion and overall spatial mapping fidelity^8^. To maximize transcriptome coverage and preserve spatial fidelity, we extensively optimized fixation, FFPE processing, and embedding to maintain high RNA integrity and tissue morphology. Despite the use of positively charged slides and optimized sectioning protocols, scalp sections, comprising tissue compartments with markedly different viscoelastic properties, frequently exhibited partial detachment and micro-drift during processing, leading to local distortion of tissue architecture and increased risk of transcript diffusion. We therefore adopted hydrogel-coated slides that covalently immobilize primary amine groups, improving tissue adherence and preserving *in situ* RNA localization (see Methods). Collectively, these optimizations achieved ultra-high transcript capture and robust spatial preservation in our Visium HD data (see below).

For spatial analyses, we retained 717,767 8µm bins with at least 50 genes/bin from four healthy adult scalp samples after tissue registration, spatial alignment, and quality control (Fig. 1d). Visual inspection confirmed that retained bins overlapped cellular regions, whereas acellular regions containing mostly collagen and subcutaneous adipose were largely excluded due to low gene detection (Extended Data Fig. 1d). Consistent with cell type-dependent capture characteristics observed in our SC datasets (Extended Data Fig. 1e), epithelial bins showed the highest transcript recovery (median 360 genes and 457 UMIs per 8µm bin across all keratinocytes), with particularly deep capture in the hair matrix (median 655 genes and 1,096 UMIs; max 5,096 genes and 11,759 UMIs per 8µm bin) (Extended Data Fig. 1f). These data underscore the high transcriptional activity and cell density in the hair matrix during anagen growth (Extended Data Fig. 1g).

Taking advantage of the micron-scale resolution and deep coverage of the transcriptome in our spatial data, we constructed a joint embedding that integrated single-cell transcriptomics datasets (both scRNA-seq and snRNA-seq) with spatial transcriptomic bins. Using TACCO, an optimal transport-based framework for integrative single-cell and spatial transcriptomics analyses^15^, we mapped single cell cluster annotations onto ST tissue sections to produce a paired spatial atlas. TACCO computed per-cluster probabilities for each bin based on gene expression and assigns each bin the highest probability annotation. Although each 8µm bin is not cell segmented and can contain mixed transcriptomes from adjacent cell types, the underlying cell heterogeneity and tissue organization was retained in the full cluster-probability matrix for downstream analyses. This spatial mapping, supported by hundreds to thousands of detected transcripts per bin, distinguished closely related sub-populations and permitted precise reconstruction of keratinocyte and stromal subtypes in their native tissue locations (Fig. 1e-f and Extended Data Fig. 1h-i), facilitating identification of some previously elusive cell populations that are difficult to resolve by markers alone but become evident when interpreted in their canonical spatial context.

The resulting spatially informed annotations recovered keratinocyte identities along the IFE-HF axis, with relative positions reflecting the layered organization of the follicular unit (Fig. 1d). Consistently, the SC UMAP and higher resolution clustering revealed an interconnected epithelial topology that extends from the IFE (*KRT14+, SOX9-*) to infundibulum (*MOXD1+, S100A8+*), through the sebaceous gland (*PPARG+, SCD1+*) and hair follicle stem cell (HF-SC) region (*CD200+, KRT15+*), extending into the lower hair follicle *(COMP+, SOX9+)*, and ending in the hair bulb/matrix (*MSX2+, LEF1+*), mirroring the physical organization of the HF unit (Fig. 1c and 1g). A similar epithelial topology was observed in the ST embedding (Fig. 1d). This correspondence was more intuitive because contiguous 8µm bins were not constrained by cell borders and therefore often captured mixed transcriptomes from adjacent cell types. Such “mixture bins” served as transcriptome-level bridges that co-localize neighboring compartments in low-dimensional embeddings. This effect was particularly evident in the bulb, where DP cells, hair shaft (HS) cortex cells, and melanocytes, populations with distinct lineages and divergent transcriptomes, clustered together in the ST embedding due to their close physical proximity within the matrix (Fig. 1d, circled). Importantly, integrating these spatial profiles with our SC reference and performing probabilistic mapping deconvolves this local mixing, enabling accurate assignment of closely apposed cell identities and resolving these entangled populations (see below). Together, integration of the SC and high-resolution ST datasets establishes a spatially resolved reference atlas of human scalp that captures both broad cell composition and fine-grained tissue organization, permitting high-fidelity cell type assignment and downstream lineage- and disease-focused analyses.

### Impact of Gene Capture on Mapping Fidelity

To examine the fidelity of our spatial mapping, we next benchmarked mapping performance in the interfollicular epidermis (IFE), which provides an ideal internal standard owing to its densely packed cell layers, stereotypical stratification and well-defined transcriptional gradients. Higher-resolution subclustering of IFE keratinocytes identified a single basal cluster and three suprabasal clusters corresponding to spinous, early granular and late granular layers (Extended Data Fig. 2a). SC-ST mapping recovered the expected layered architecture toward the stratum corneum, evident both in categorical bin assignments and in layer probability heatmaps (Extended Data Fig. 2b–d). This computationally mapped stratification was corroborated by canonical spatial marker patterns, including KRT5/KRT14 (basal), KRT1/DSG1 (spinous), KRT2/KLK7 (early granular) and IVL/ALOX3 (late granular), supporting the effective resolution and accuracy of our spatial mapping strategy (Extended Data Fig. 2d).

In the densely packed HF bulb/matrix, closely related epithelial populations are highly interleaved, and this creates an additional challenge for spatial mapping. In the matrix epithelial compartment, our optimized assay detected 17,954 expressed genes. Importantly, ultra-high transcriptome capture per bin was particularly critical for resolving nuanced identities, enabling clear delineation of matrix-associated layers and faithful mapping of these identities to their known anatomical location, 3-5 mm below the epidermis (Extended Data Fig. 3a). Although Visium HD does not incorporate explicit cell segmentation (in contrast to imaging-based platforms such as Xenium), we found that reducing the gene set to 5,000 genes, either by random subsampling or by restricting to the Xenium Prime 5K Human panel overlap (4,807 of 5,001 genes), decreased layer clarity and increased mis-localization of matrix-associated identities to epidermal and upper-HF regions in both scenarios (Extended Data Fig. 3a). Further reductions to 3,000–500 genes produced progressive deterioration in SC–ST mapping and increased spatial bin mis-assignments (Extended Data Fig. 3a), indicating that high-dimensional transcriptomic anchors are required to match the complexity of scalp tissue architecture and to distinguish closely related states *in situ*.

To quantify how spatial gene capture governs mapping fidelity, we assessed the effect of RNA integrity on Visium HD performance. Among four Visium HD libraries, three were derived from samples with high-quality RNA (DV200 >80%) and one with lower-quality RNA (DV200 ∼40%). Although all samples exceeded the recommended DV200 threshold (>30%), the DV200 ∼40% sample yielded markedly fewer UMIs and genes per 8 µm bin (Extended Data Fig. 3b). While its output remained broadly comparable to previously reported spatial transcriptomics datasets, this reduction in capture substantially degraded the fidelity of SC-ST integration and fine-resolution mapping (Extended Data Fig. 3c-d). In the lower-quality sample, epidermal layering became discontinuous and bins overlying stromal compartments, such as fibroblasts, were frequently filtered out because of reduced gene detection (Extended Data Fig. 3c). A similar loss of keratinocyte layering and stromal coverage was observed in the hair matrix, a region that exhibited among the highest transcript abundance across the scalp (Extended Data Fig. 3d). Taken together, these analyses establish that our tissue preserving workflow, high RNA quality (DV200 >80%) and ultra-high transcript capture in micron-scale bins, enabled by our optimization, are critical for accurate integration of single-cell and spatial datasets and for faithful reconstruction of tissue architecture.

### Spatially-Restricted Identification of Hair Follicle Stem Cells

The hair follicle (HF) harbors a population of label-retaining epidermal stem cells that is essential for cyclic HF growth and contributes to epidermal repair after tissue injury^16^. These cells, often called hair follicle stem cells (HF-SCs), are located in the “bulge” region in mice, positioned below the sebaceous gland (SG) and named for the outward curvature of this epithelial compartment^17–20^. However, HF morphology and cycling behaviors differ substantially between mice and humans. In particular, the human anagen HF lacks a distinctive bulge morphology. Furthermore, whereas ∼90% of adult murine follicles rest in quiescent telogen^21,22^, ∼90% of human scalp follicles persist in anagen, with growth phases lasting ∼3-10 years. Given the central importance of HF-SCs in HF maintenance and epidermal regeneration, high-resolution interrogation of human HF-SCs and their niche *in situ* is critical.

Our single-cell atlas identified a transcriptomically distinct basal keratinocyte population topographically situated at the junction between basal ORS and SG keratinocytes in the UMAP embedding, raising the possibility that this population may represent human HF-SCs. Spatial mapping localized these putative HF-SCs to the basal HF sheath just below the SG and abutting the arrector pili muscle (APM) (Fig. 2a). DEG analysis comparing the HF-SC cluster with all IFE and HF keratinocytes uncovered a comprehensive HF-SC gene signature (Supplementary Table 3). Genes associated with HF keratinocyte differentiation, including structural components and transcription factor (TF) regulators, such as *TCHH, KRT27, KRT83, DSG1, DSG4, FOXN1, GRHL3,* and *ZNF750*, were among the most depleted, underscoring an undifferentiated cell state. Interestingly, *FGF5*, a gene linked to human HF regression^23^, was also among the most depleted signaling genes in the anagen HF-SC population.

**Fig. 2:**
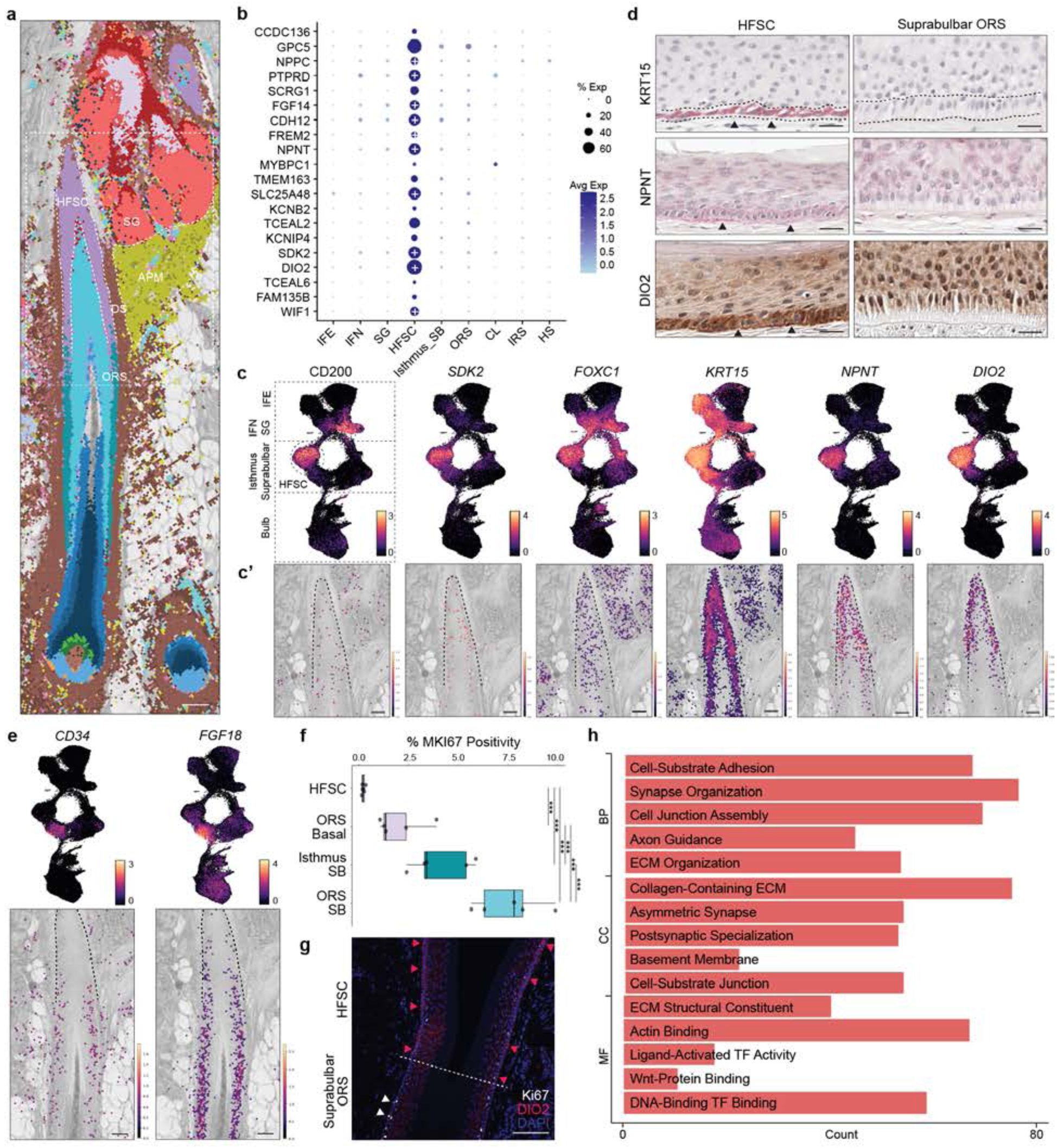
Identification of Human HF-SCs *in situ*. **(a)** Visualization of TACCO-mapped single-cell cluster identities for a representative anagen follicular unit with grayscale H&E overlay. SG: sebaceous gland. APM: arrector pili muscle. DS: dermal sheath. ORS: outer root sheath. Scale bar: 100µm. **(b)** Dot plot of HF-SC marker genes compared to other keratinocyte populations. Plus sign (+) denotes gene being detected as differentially expressed for HF-SCs in ST data as well. **(c-c’)** UMAP feature plot (c) and spatial gene expression visualization (c’) of the HF-SC region for canonical and novel HF-SC markers. Circled: HF-SC cluster on single-cell atlas UMAP. Dashed line: HF-SC region. Scale bar: 100µm. **(d)** IHC of HF-SC markers demonstrating HF-SC specific expression (arrowheads) in the basal keratinocytes in the isthmus region compared to suprabulbar ORS cells. Scale bar: 100µm. **(e)** UMAP feature plot and spatial gene expression patterns for mouse HF-SC markers *CD34* and *FGF18* showing minimal expression in the human HF-SC region and a downward shift to the ORS. **(f)** Box plot of the percentage of cells within each ORS subpopulation showing *MKI67* RNA expression. ***denotes statistical significance from pairwise contrasts of estimated marginal means. **(g)** Ki67 and DIO2 staining of the HF-SC region showing no basal Ki67 expression in the DIO2+ HF-SCs (red arrowheads) compared to the basal (white arrowheads) and suprabasal Ki67 positivity in the ORS. Dashed line: HFSC-ORS demarcation by cell morphology and DIO2 expression. **(h)** Gene Ontology biological process (GO: BP), cellular component (GO: CC), and molecular function (GO: MF) terms derived from differentially expressed marker genes comparing HF-SCs with other basal keratinocyte populations.

Next, we examined the expression patterns of key marker genes across single-cell clusters and within the spatially-defined HF-SC region. These analyses confirmed the expression of established HF-SC markers, such as *KRT15*^24^,*CD200*^25,26^, and *DIO2*^27,28^, and identified a plethora of highly specific candidates such as *NPNT*, *TMEM163, WIF1, DCN, FGF14,* and *SDK2* (Fig. 2b-c and Supplementary Table 3). Immunohistochemistry (IHC) staining for selected markers, including KRT15, NPNT and DIO2, validated their specificity in the HF-SC region relative to immediately adjacent ORS cells (Fig. 2d and Extended Data Fig. 4a). In contrast, canonical bulge markers in mice, including *CD34* and *FGF18*^18,19,29^, were depleted in human HF-SCs and instead shifted downward to a more distal ORS compartment, highlighting the species divergence (Fig. 2e). Conversely, several functional regulators, which are experimentally linked to HF-SC adhesion and tissue organization in mouse models, such as *FOXC1*^22,30,31^, *LHX2*^32^, and *NPNT* ^33^, appeared conserved in human scalp HF-SCs (Fig. 2c-d, Extended Data Fig. 4b and Supplementary Table 4). Notably, many WNT antagonists, including *WIF1, DKK3, SOSTDC1,* and *SFRP1*, were enriched in HF-SCs, indicating a locally WNT-inhibitory microenvironment (Extended Data Fig. 4c and Supplementary Table 4). Quiescence-associated factors described in murine HF-SCs, including *BMP6* and *FGF18*, were more highly expressed in ORS progenitors than in human HF-SCs, mirroring the pattern of *CD34* (Supplementary Table 4). Nevertheless, analysis of Ki67 mRNA levels suggested that the HF-SC compartment is less proliferative and more quiescent than basal ORS cells, indicating a quiescent HF-SC state during prolonged anagen growth (Fig. 2f). Consistently, Ki67 immunostaining revealed that Ki67+ basal keratinocytes were rare within the HF-SC region (DIO2+) compared to areas immediately above and below along the HF sheath (Fig. 2g and Extended Data Fig. 4d).

Interestingly, thyroid-hormone metabolism exhibited a striking polarity within the HF sheath: DIO2, which activates thyroid-hormone (TH) signaling by converting inactive pro-hormone thyroxine (T4) to the active triiodothyronine (T3), was among the most enriched HF-SC genes (Fig. 2c-d), whereas DIO3, which converts active T3 to inactive T2, was strongly depleted (Supplementary Table 4). This *DIO2*-high/*DIO3*-low signature predicts elevated TH signaling in the anagen HF-SC compartment, consistent with recent data linking topical T3 to anagen hair growth in humans^34^. Probing more broadly, Gene Ontology (GO) analysis of HF-SC-enriched genes further highlighted ECM organization, cell-substrate adhesion, cell junction and transcriptional regulation as key features of the HF-SC transcriptome (Fig. 2h). Collectively, our atlas reveals human HF-SCs as a spatially restricted, undifferentiated and quiescent basal keratinocyte population during anagen. The identification of conserved and human-specific markers provides a molecular framework for selective purification, based on cell surface markers (Extended Data Fig. 4e) and functional interrogation of this compartment in future studies.

### Spatial Mapping of Heterogeneous Stromal Subclusters

Classical histology is founded on the principle that tissue function is intimately tied to the spatial arrangement of its constituent cells^35^. Dissociation-based scRNA-seq digitizes cellular states into high-dimensional data at scale, yet the assignment of clusters is highly sensitive to computational parameter choice and batch effect. The problem is particularly acute for closely related stromal populations, such as the dermal fibroblast populations of the skin^36,37^. Without independent spatial validation, it is unclear whether computationally defined clusters represent *bona fide* cell populations or analytic artifacts. To determine the identities of stromal populations, including highly heterogeneous fibroblast subpopulations, from our SC dataset and further inform tissue architecture, we performed subclustering and spatial mapping of fibroblast and related smooth muscle populations. The higher resolution subclustering revealed 6 fibroblast populations, with the global atlas-level Fib I, II, and III populations each giving rise to 2 subpopulations. Smooth muscle populations, on the other hand, stably maintained the same clustering pattern as SM I, SM II, and APM, marked by shared *ACTA2* expression (Fig. 3a-b).

**Fig. 3:**
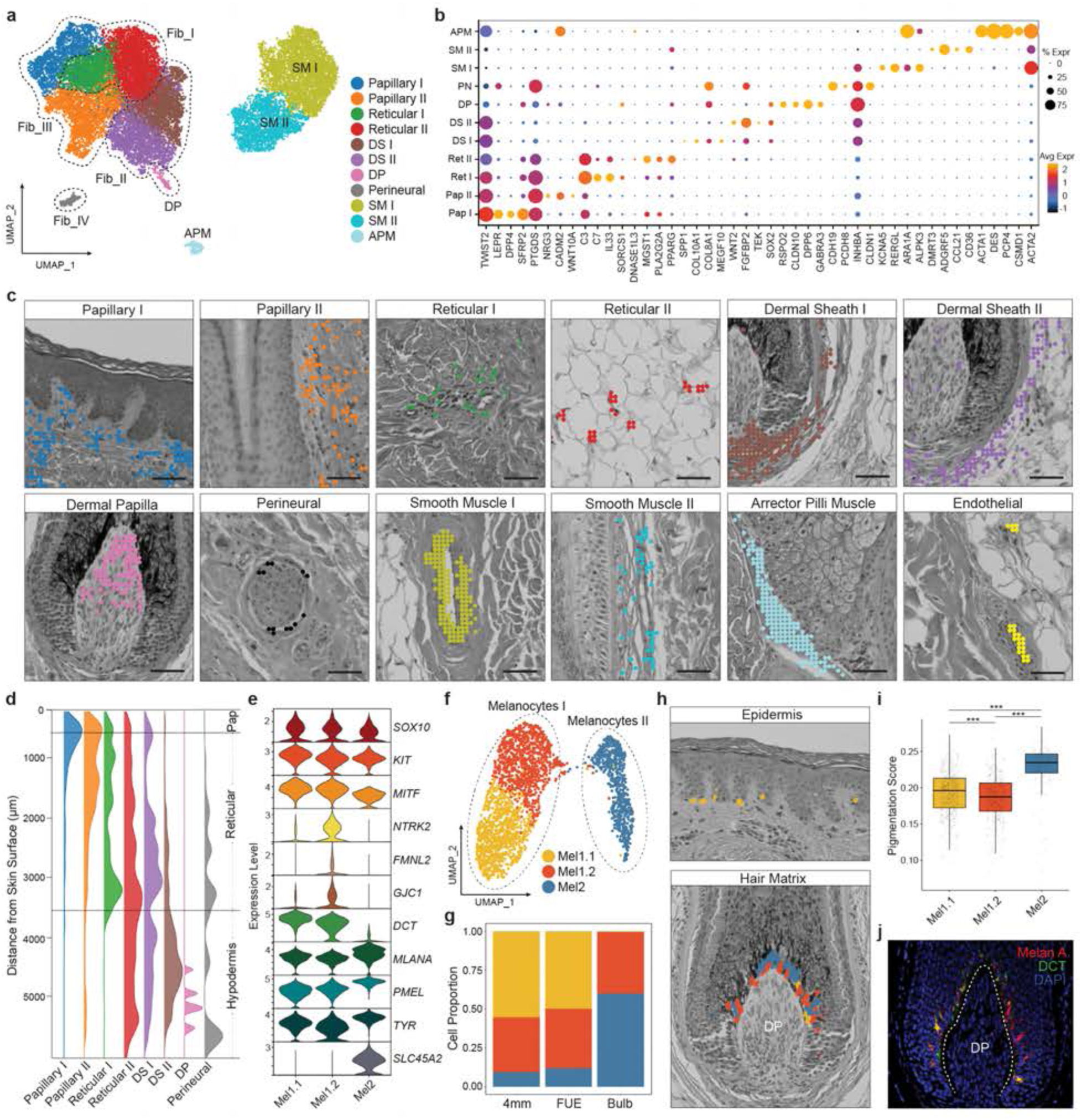
Spatial Deconvolution of Fibroblast and Melanocyte Subtypes. **(a)** UMAP of high-resolution subclustering of fibroblasts (Fib I, II, III, VI, DP) and smooth muscle (SM I, SM II, APM) populations identified from atlas-level annotations. DS: dermal sheath. DP: dermal papilla. SM: smooth muscle. APM: arrector pili muscle. **(b)** Dot plot of marker gene expression for fibroblast and smooth muscle subtypes. **(c)** Visualization of spatial mapping results for fibroblast and smooth muscle subpopulations with H&E overlay. Scale bar: 50µm. **(d)** Ridgeline plot of the spatial density distribution of fibroblast subtypes along the epidermis-hypodermis (superficial-deep) axis. **(e)** Violin plot of selected melanocyte subpopulation markers and melanocyte differentiation genes. **(f)** UMAP plot of melanocyte subpopulations. Subpopulations are annotated based on population of origin within the global clustering. **(g)** Relative proportions of each melanocyte subpopulation within the three scRNA-seq isolation methods showing no contribution of Mel1.1 in the hair bulb mechanical isolation samples. **(h)** Spatial mapping of melanocyte subpopulations within the interfollicular epidermis (top) and hair matrix (bottom) using cell-segmented spatial data. Plotted points represent 2µm bins labeled by their melanocyte annotations. **(i)** Box plot of pigmentation gene module scores within each melanocyte subpopulation. ***denotes statistical significance from two-tailed Mann-Whitney U test. **(j)** Melan-A and DCT staining of the hair matrix showing Melan-A-high; DCT-low as well as double-positive melanocytes in the hair matrix corroborating the Mel1.2 and Mel2 spatial mapping.

To determine biological relevance of these finely clustered stromal cells, we mapped these cell clusters onto our ST dataset using TACCO to facilitate annotation transfer between SC and our deep-coverage spatial data^15^, and visualized their spatial distribution patterns across the skin. Strikingly, each of these 6 stromal subpopulations mapped to distinct spatial locations, which helped reveal their identities and function. One of the Fib III subclusters mapped to a papillary fibroblast population (Papillary I), which was spatially distributed in the immediate layer below the dermal-epidermal junction (DEJ) of the epidermis, marked by *TWIST2, LEPR*, and *DPP4* (Fig. 3b-d, Extended Data Fig. 5a and Supplementary Table 5). Although Papillary II was also found along the DEJ of the IFE, they were located primarily along the DEJ of the upper HF infundibulum and the SG until a transition to dermal sheath (DS) fibroblast populations below the SG (Fig. 3c-d and Extended Data Fig. 5a). Thus, our single-cell spatial mapping results further distinguished a papillary fibroblast population, located in the upper HF (uHF) regions, forming a sheath-like structure with potential roles in establishing a niche for uHF keratinocytes.

In contrast, Fib I subclusters mapped to two populations in the reticular dermis, with Reticular I primarily localizing around blood vessels at the dermal-hypodermal junction and Reticular II interspersed in the dermal white adipose tissue (dWAT) (Fig. 3c-d and Extended Data Fig. 5b). Examination of marker gene expression of Reticular I cells (Fig. 3b and Supplementary Table 6) revealed enrichment for immune regulatory functions, including chemokine production, lymphocyte differentiation and regulation of inflammatory response (Extended Data Fig. 5c), consistent with their spatial location predominantly surrounding vasculature. Conversely, Reticular II fibroblasts exhibited enrichment for regulation of cholesterol transport and fibroblast proliferation (Fig. 3c, Extended Data Fig. 5c and Supplementary Table 6), as evidenced by upregulation of *PPARG*, *MGST1*, and other adipogenic transcription signatures. Thus, these enriched gene expression programs were in concordance with their spatial localization in dWAT and revealed adipogenic identity of Reticular II fibroblasts.

Fib II subclusters, renamed to dermal sheath I (DS I) and II (DS II) upon subclustering, mapped to two lower perifollicular fibroblast populations, traditionally considered as the dermal sheath. Notably, our spatial mapping uncovered a bi-layered split in the DS population, which has not been previously described (Fig. 3c and Extended Data Fig. 5a), likely owing to the lack of knowledge for their spatial and transcriptomic identities. DS I was proximal to the HF ORS and responsible for the deposition of ECM into the perifollicular region, corresponding to its higher expression of *COL8A1* and *COL10A1* (Fig. 3b). Gene programs associated with TGFβ signaling and smooth muscle contraction were also highly enriched in DS I (Extended Data Fig. 5c and Supplementary Table 7), likely reflecting its potential contribution to catagen progression through concentric contractions^38^. DS II, on the other hand, was more net-like in nature and supported a complex neurovascular plexus, consistent with the enrichment for nerve development and VEGFR signaling (Extended Data Fig. 5c and Supplementary Table 7). DS II was noted to express higher levels of *WNT2* and *TEK* genes, which have been previously described to play important roles in mechanosensing and modulation of angiogenesis^39–41^. Interestingly, although DS I was more proximal to the lower HF regions, such as the lower proximal cup (LPC) and ORS, some DS II cells mapped to the lower DP area (Fig. 3c), suggesting a potential association with the DP, which is a highly specialized fibroblast population and functions to regulate HF growth^42,43^.

Because human DP has been poorly captured in previous studies, we performed the spatial mapping and in-depth analysis of gene expression with our SC and ST datasets. Notably, the specialized DP population, well-identified in both the global atlas (Fig. 1c-d) and fibroblast subclustering (Fig. 3a-b), robustly mapped to the DP location in the lower HF bulb and marked by canonical marker genes identified in mouse studies^44–46^, including *RSPO2, RSPO3, CORIN, CRABP1*, *HHIP* and *SCUBE3* (Fig. 3b-c and Extended Data Fig. 5d). Leveraging the deep transcriptome coverage of our datasets, we identified genes that are enriched in the DP. Notably, WNT signaling, represented by multiple layers of regulators such as *RSPO2/3/4, WLS, WNT5A, DKK3, SOSTDC1* and *LEF1* (Extended Data Fig. 5d and Supplementary Table 5), was highly enriched in the DP. *BMP5* and *BMP7* were also highly enriched in the DP whereas *BMP6* was more enriched in the DS II (Extended Data Fig. 5d). *FGF10*, known to stimulate hair growth in mouse^47^, was also enriched in the DP (Extended Data Fig. 5d). Interestingly, some genes showed differential proximal vs. distal expression patterns, such as *EDN3* and *RSPO2,* whereas other genes, such as *DIO3* and *MGP,* showed more pan-DP expression (Extended Data Fig. 5e), revealing spatially defined heterogeneity in the DP. This differential pattern, in turn, led to mapping of many DP cells to the upper DP, whereas the lower DP was occupied by some DS cells instead (Fig. 3c and Extended Data Fig. 5a). This spatial arrangement of the DP and DS II cells could reflect a putative lineage relationship of HF dermal stem cells (hfDSC) to the DP^48^.

Fib IV, another specialized fibroblast population, mapped exclusively to the perineural sheath of nerves in the skin, marked by *CDH19, CLDN1*, and *PCDH8* (Fig. 3a-c). This fibroblast population with the enrichment for myelination and receptor localization to synapse (Extended Data Fig. 5c) was distinct from Schwann cells, which were found intercalated within the nerve bundles. In support of this notion, canonical Schwann cell markers *SOX10, MPZ,* and *MBP* were absent in these perineural fibroblasts (Extended Data Fig. 5f). For smooth muscle populations, the APM, expressing high levels of *ACTA2, DES, PCP4*, and *ACTG2* (Supplementary Table 5), reliably mapped to bundles of smooth muscle in the upper dermis, some coursing around the SGs in the HF isthmus (Fig. 3b-c). SM I mapped to the muscular walls of larger arterioles/arteries. In conjunction with their canonical marker expression, such as *TAGLN, CNN1, KCNA5*, and *PRDM16*, these cells were identified as contractile vascular smooth-muscle cells (Fig. 3b-c). SM II dotted the perimeters of smaller vessels, frequently adjacent but external to endothelial cells and most densely distributed in the outer dermal sheath plexus of the lower HF. These cells express *RGS4, KCNJ8, ADGRF5, ANGPT2* and other canonical pericyte markers, while being negative for endothelial markers such as *PECAM1, CDH5,* and *VWF* (Extended Data Fig. 5g).

Finally, we leveraged spatial mapping to resolve melanocyte heterogeneity across epidermal and follicular compartments. Two melanocyte populations (Mel I and Mel II) emerged from the global atlas (Fig. 1c and 1g), and higher-resolution subclustering further resolved three transcriptionally distinct melanocyte states, hereby termed Mel 1.1, 1.2, and 2 reflecting their initial cluster of origin (Fig. 3e-f). Mel 1.1 was absent from mechanically dissected bulb preparations, consistent with an epidermal origin, whereas Mel 1.2 and Mel 2 were enriched in bulb samples (Fig. 3g). Spatial mapping confirmed these inferenced distributions, localizing Mel 1.1 to IFE and Mel 1.2/2 to the hair matrix (Fig. 3h and Extended Data Fig. 5h). DEG analysis indicated relatively subtle differences between Mel 1.1 and Mel 1.2, whereas Mel 2 showed higher expression of melanogenesis-associated genes (Fig. 3e). To quantify melanogenic potential, we computed a pigmentation score using a curated panel of 169 pro-melanogenic genes identified in a genome-wide CRISPR screen^49^. Mel 1.1 exhibited an intermediate pigmentation score, while the two matrical populations diverged, with Mel 1.2 displaying low and Mel 2 high pigmentation scores, respectively (Fig. 3i). In addition, Mel 2 expressed elevated melanocyte differentiation markers, such as *MLANA, PMEL, TYR*, *TYRP1,* and *SCL45A2*, a solute carrier regulating melanosome pH balance during late melanogenesis^50,51^, further supporting its characterization as a fully differentiated melanosome producer responsible for hair shaft pigmentation (Fig. 3e). Consistent with this melanocyte cell organization inferred from spatial mapping, IF for DCT and MLANA revealed both DCT^Hi^ and DCT^Lo^/Melan-A+ melanocytes within the matrix, in accordance with the spatial mapping and inferred expression patterns (Fig. 3j). Although both SC and ST analyses did not resolve a discrete melanocyte stem cell (MSC) cluster, possibly due to scarcity, we detected rare SOX10+ cells in HF-SC/ORS regions and more abundantly in the matrix, consistent with the localization of Mel 1.2 and Mel 2 populations (Extended Data Fig. 5i-i’). Taken together, joint spatial mapping of the SC and deep-coverage ST datasets provides an unprecedented framework to elucidate stromal compartments, assign anatomically grounded identities to closely related fibroblast populations, and resolve melanocyte states across epidermis and follicular microenvironment, thereby enabling spatially interpretable hypotheses about cellular heterogeneity and lineage relationships.

### Joint Contribution of Two Cell Lineages for Sebaceous Gland Formation

Thus far we have relied on paired SC and ST datasets to validate cell type assignments and discover tissue architecture. However, this strategy is challenging for the SG because human sebocytes are large, lipid-laden cells that are rarely captured by droplet-based scRNA-seq platforms, even less so than their murine counterparts^52^. Indeed, our SC atlas was skewed toward SG-related ductal keratinocytes, marked by basal *KRT5/COL17A1+* and suprabasal *S100A9/KRT6A+* keratinocytes, respectively (Extended Data Fig. 6a-b). In contrast, only a much smaller population of *KRT5/PPARG+* basal SG progenitors and a handful of terminal sebocytes, expressing *MGST1, KRT79, CIDEA, AWAT2* and *FA2H* ^53,54^, were recovered (Extended Data Fig. 6a-b), despite their abundance in intact scalp skin (Fig. 4a and Extended Data Fig. 6c).

**Fig. 4:**
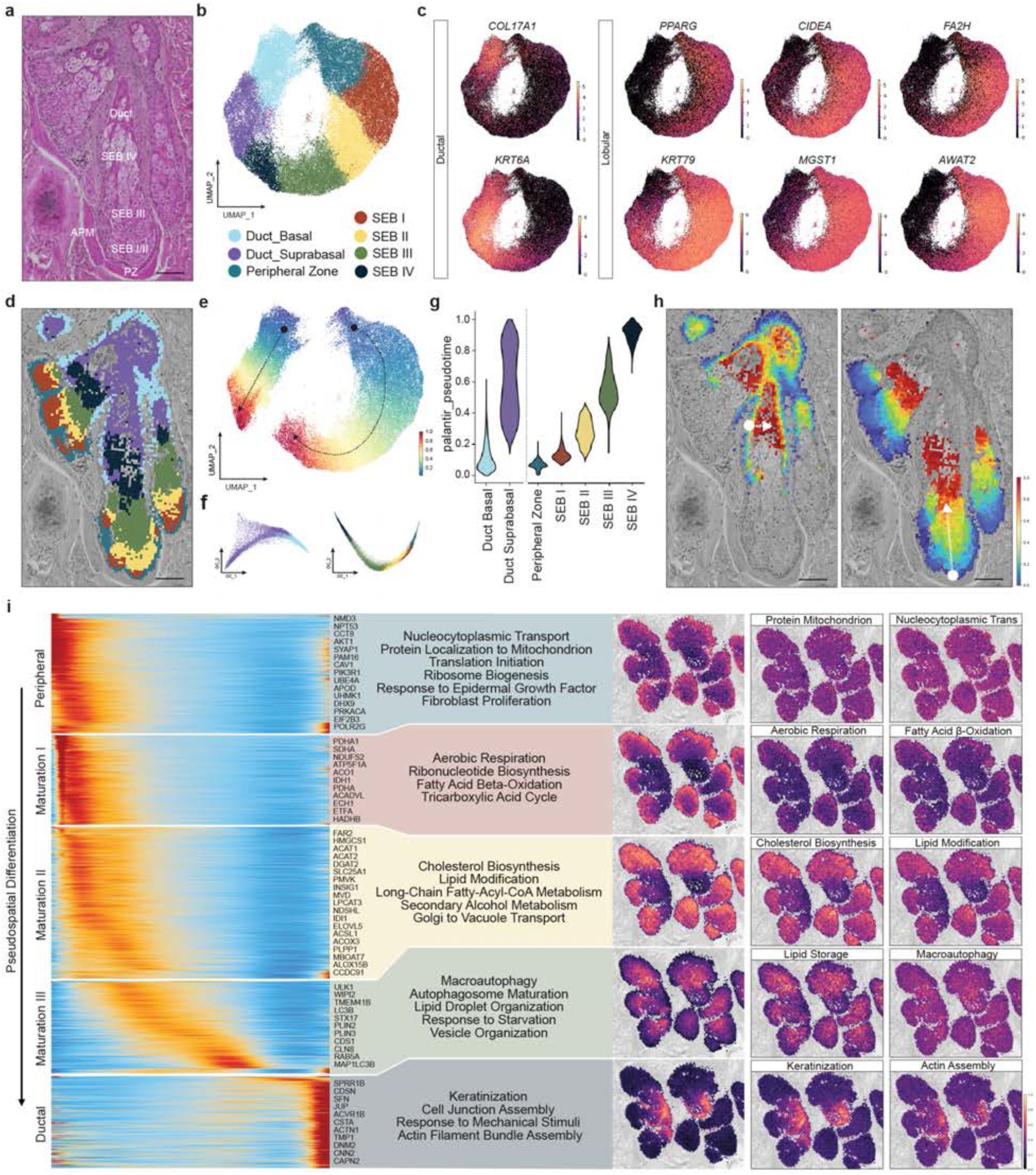
Spatial Transcriptomic Analysis of Human Sebaceous Gland *In Situ*. **(a)** H&E image of a representative human sebaceous gland and associated ductal structures. APM: arrector pili muscle. PZ: peripheral zone. SEB: sebocyte. Scale bar: 100µm. **(b)** UMAP of high-resolution subclustering of ST 8µm bins associated with the sebaceous gland. **(c)** UMAP feature plots for canonical sebaceous gland Ductal (*COL17A1, KRT6A*) and Lobular (*PPARG, KRT79, CIDEA, MGST1, AWAT2, FA2H*) genes. **(d)** Visualization of high-resolution SG subclusters with H&E overlay. Scale bar: 100µm. **(e-f)** Palantir pseudotime heatmaps of the Ductal (left) and Lobular (right) trajectories (e) and their associated diffusion maps (f). **(g)** Violin plot of the ranked Palantir pseudotime values for the Ductal and Lobular trajectories. **(h)** Visualization of the projected Palantir pseudotime values for the Ductal (left) and Lobular (right) trajectories with H&E overlay. Scale bar: 100µm. **(i)** Pseudotime heatmap of Lobular trajectory genes grouped into five modules (left). Representative genes and GO terms associated with the pseudospatial modules (middle). Visualization of the collective pseudospatial module and select individual biological processes (GO: BP) with H&E overlay (right).

To address this issue, we leveraged the micron-scale resolution and deep coverage of our ST datasets alone to reconstruct SG lineage trajectory and their spatial organization *in situ*, independent from the scRNA-seq data. Across four scalp ST samples, we obtained 56,104 8µm bins that identified the sebaceous duct/gland regions by Louvain clustering (Extended Data Fig. 6c). Across these 8µm spatial bins, we captured a median of 583 UMI (min = 154, max = 5,142) and 467 genes per bin (min = 151, max = 2,488), which readily outperformed the recently published results from Visium HD and Stereo-seq^8^. Strikingly, in the UMAP embedding, these bins formed an annular arrangement with one arm (left) enriched for *COL17A1/KRT6A,* identifying ductal keratinocytes, and the other enriched for *PPARG, KRT79, CIDEA, MGST1, FA2H,* and *AWAT2,* characteristic of sebocytes that were scarcely recovered in our SC data (Fig. 4b-c). Visualization of the SG subclusters overlaid on H&E images revealed that the Peripheral Zone (PZ, cyan) progenitors localized at the distal perimeter of the SG lobules (Fig. 4d) while differentiating sebocytes (SEB I-IV) were arranged centripetally toward the center of the SG lobule (Fig. 4d). Conversely, the sebaceous duct consisted of a basal layer (Duct_Basal, light blue) and a multilayered suprabasal compartment (Duct_Suprabasal, purple) (Fig. 4d). Terminally differentiated sebocytes (SEB IV), while egressing from the gland, closely interdigitated with ductal suprabasal cells. Notably, the spatial bins (count = 3,418 representing 6% of SG area) that located in this area captured this intermixed interface (Fig. 4b and 4d, dark green), as evidenced by both their location and their mixed transcriptome, characteristic of both terminal sebocytes and ductal suprabasal keratinocytes. These intermixed bins created a bridge on the UMAP embedding, effectively connecting the ductal and sebocyte lineages into a continuous loop (Fig. 4b-c). Such mixed-transcriptome bins, generated where different cell types are in consistent and intimate contact, encoded informative spatial relationships of distinct cell types, a phenomenon also seen in the global ST UMAP topology where the DP, hair shaft cortex, and hair matrix melanocytes were bridged together due to their physical association in the hair bulb (Fig. 1d, circled)

Because the spatial bins segregated the SG cells into two developmental trajectories, we computed their pseudotime separately for each lineage with the earliest pseudotime values rooted at the basal layers (Duct_Basal and PZ), respectively (Fig. 4e-g). Ductal keratinocytes matured linearly along the basal-suprabasal axis, whereas sebocytes matured concentrically from the outer PZ through SEB I-III, culminating in fully mature SEB IV sebocyte cells that discharge their contents into the SG duct for sebum excretion (Fig. 4h). In mice, lineage tracing studies showed that ductal and sebocyte lineages are maintained independently^55^. However, there is no prominent ductal suprabasal population in mice, and sebocytes differentiated linearly from the SG lobule base toward the hair shaft canal^52^. To validate these ST-based findings, we performed IHC analysis of COL17A1 and KRT6 expression patterns. In support of our ST data, the sebaceous duct is composed of a single COL17A1+ basal layer and multiple KRT6+ suprabasal keratinocyte layers with highly eosinophilic, thin keratin projections into the ductal space, leading to interdigitation with terminal sebocytes as predicted by the mixture bins of our ST SG dataset (Extended Data Fig. 6d-e). Therefore, the radial lobular pattern and expanded ductal compartment we observed in human scalp likely represent structural adaptations to accommodate the enlarged gland size and enhanced secretory function.

With the SG lobular pseudotime computed, we further dissected the differentiation programming of the SG lobule and visualized these programs on the SG to reveal where these events occurred *in situ*. Among 17,898 detected genes whose expression spans nearly three orders of magnitude in raw expression matrix, we identified the 1,500 most dynamic genes along the SG lobular pseudotime trajectory and performed Louvain clustering by their expression dynamics. This revealed 5 spatially resolved modules: Peripheral, Maturation I-III, and Ductal (Fig. 4i left panel and Supplementary Table 8). We then performed GO analysis on these gene modules to determine the activated molecular pathways along the SG lobular cell lineage *in situ* (Fig. 4i middle panel). When these modules were mapped onto their original spatial bins, it revealed the Peripheral module primarily corresponded to PZ progenitor cells located in the basal layer with enrichment for cell proliferation, protein synthesis, and mitochondrial activation (Fig. 4i right panel and Extended Data Fig. 6f). Subsequently, Maturation I encompassed the PZ and the first 2-3 lobular cell layers where an increase in energy utilization, TCA cycle, fatty acid β-oxidation was observed, creating the building blocks required for sebum production (Fig. 4i right panel and Extended Data Fig. 6f). The next 2-3 sebocyte layers were represented by the Maturation II program enriched for cholesterol and lipid synthesis and modifications. These functions were in accordance with studies of sebum composition, containing mostly triglycerides, wax esters, squalene, and free fatty acids^56,57^. As sebocytes reached the stage of near terminal maturation (reflected by their initial entry into the SG duct), they transitioned to the Maturation III program enriched for lipid organization and packaging. In addition, macroautophagy signature was a prominent late differentiation program, consistent with the removal of cellular components as the sebocytes became laden with sebum^58,59^ (Fig. 4i right panel and Extended Data Fig. 6f). As differentiated sebocytes moved into the sebaceous duct, a gene program of keratinocyte differentiation marked by canonical suprabasal genes like *KRTDAP*, *SPINK5* and *KRT6* started to dominate in the inner ductal region, where terminal sebocytes interdigitated with ductal suprabasal cells (Fig. 4h-i and Extended Data Fig. 6g-h). Collectively, our micron-scale ST map of healthy human scalp skin overcomes the limitations of the spot size and RNA capture of recent Visium SD studies of the SG^60,61^ and delivered an unprecedented view of the human SG at near-single-cell resolution. By resolving both the concentric sebocyte lineage and the adjoining ductal trajectory and pinpointing their mixed interface, we provide a molecular and cellular blueprint of how distinct epithelial lineages cooperate to build a functional gland architecture *in situ*.

### Tissue Architecture of Hair Follicle Matrix

The hair matrix is the growth center of mammalian HFs during anagen. It has one of the highest mitotic rates in the body, coupling rapid proliferation to continuous differentiation. Matrical progenitors generate the distinct cell lineages that form the hair shaft (HS), including cortex and medulla, and inner root sheath (IRS), driving hair growth at ∼0.3-0.4mm/day in humans^10^. How this compartment is spatially organized to support the requisite growth demands remains debated. Classic models proposed a pool of highly proliferative, multipotent matrix progenitors within the hair bulb that give rise to the three HS layers and three IRS layers^62,63^. More recent mouse studies have suggested instead that lineage-restricted progenitors for each HF layer are arranged around the DP^64–66^. While these murine studies have advanced our understanding of matrical differentiation, murine pelage and human scalp hair differ substantially in growth kinetics and cycling dynamics. Adult murine HFs complete anagen growth in ∼2 weeks, with maximal hair production during anagen VI lasting less than one week, compared with years of growth in humans^12,13^. Given these prominent interspecies differences, the spatial organization and differentiation trajectories of human matrical progenitors, critical for maintaining stable tissue architecture during prolonged rapid hair growth, remain poorly defined.

In our global atlas, matrix populations formed a transcriptionally distinct cluster separate from HF-SCs and ORS progenitors (Fig. 1c). High-resolution subclustering of matrical cells revealed in a branching UMAP topology in which the base was populated by *COL17A1+* basal cells, a marker previously associated with basal epithelial cell stemness^67,68^, and the distal branches resolved the canonical IRS and HS layers (Fig. 5a). Leveraging high-resolution ST data, visualization of *COL17A1* mRNA and protein confirmed a basally located COL17A1+ layer, forming a continuous thin layer along the matrix-DP border (Fig. 5b and Extended Data Fig. 7a). Arising from this basal layer, we detected all morphologically distinct lower HF cell lineages, including the LPC (*LGR5+*), three IRS layers (Henle *KRT71+*, Huxley *KRT74+*, and IRS cuticle *KRT72+*), and three HS layers (cuticle *KRT82+*, cortex *KRT31+*, and medulla *FOXQ1+*) (Fig. 5b).

**Fig. 5:**
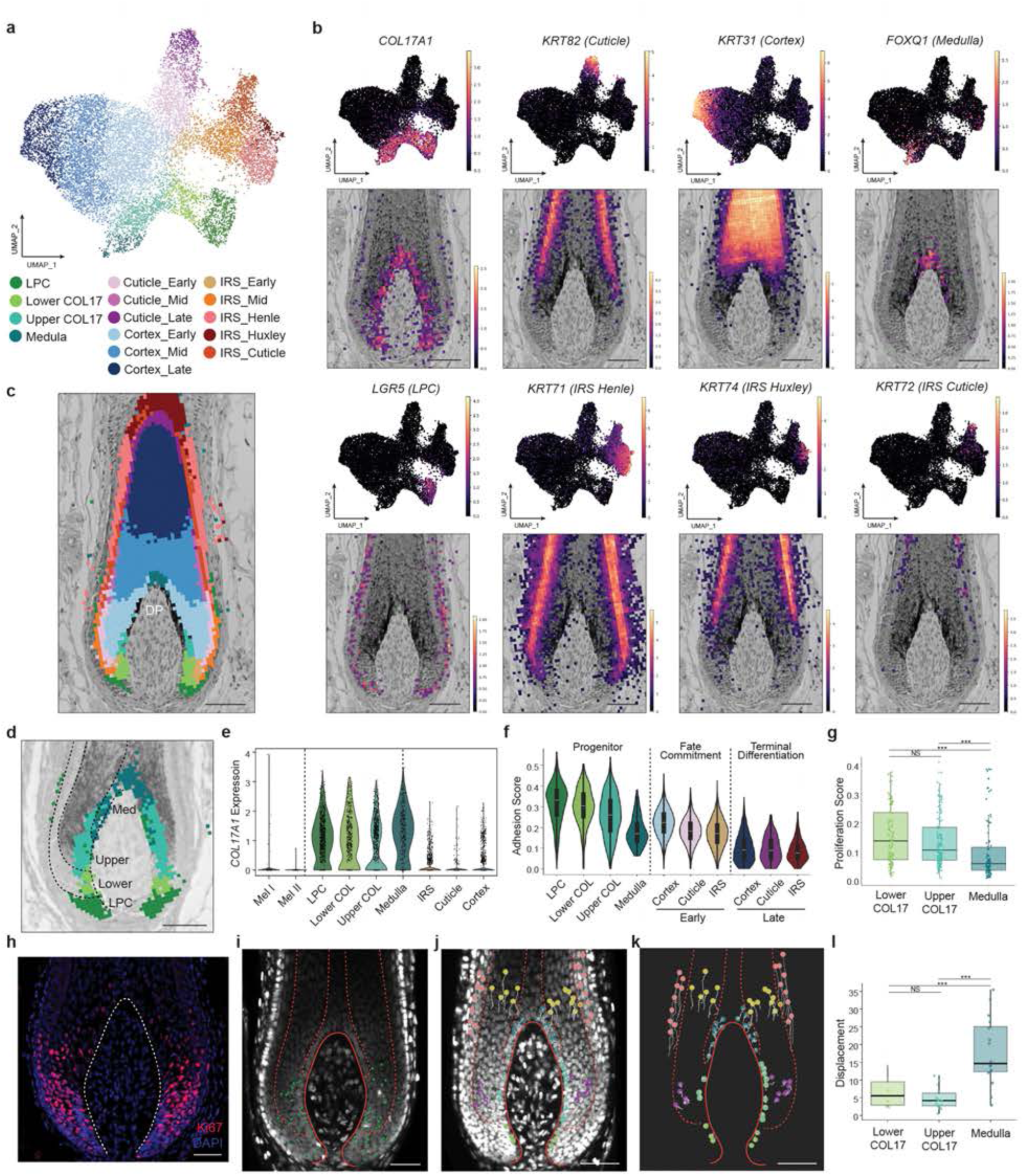
High-Resolution Spatial Mapping Reveals Tissue Architecture of Human Hair Matrix. **(a)** UMAP of high-resolution subclustering of lower hair follicle/matrical keratinocytes. **(b)** UMAP feature plots (upper panels) and spatial gene expression (lower panels) visualizations of key matrical lineage markers. Basal: *COL17A1*, lower proximal cup (LPC): *LGR5*, cuticle: *KRT82*, cortex: *KRT31*, medulla: *FOXQ1*, IRS Henle: *KRT71*, IRS Huxley: *KRT74*, and IRS cuticle: *KRT72*. Scale bar: 100µm. **(c)** Visualization of high-resolution subclustering of matrical keratinocytes with spatial overlay. Scale bar: 100µm. DP: dermal papilla. **(d)** Spatial mapping of COL17A1+ matrical populations showing clear segmentation of three populations along the matrix-DP junction. Scale bar: 100µm. **(e)** *COL17A1* expression among matrix cell populations showing high expression in matrix basal populations and minimal expression in matrix melanocytes and differentiated keratinocytes. **(f)** Violin plot of cell adhesion scores for matrical subpopulations, segmented into three groups based on differentiation: progenitors, fate commitment, and terminal differentiation. **(g)** Box plot of proliferation gene module scores across lower and upper COL17A1 progenitors and medulla progenitors. ***denotes statistical significance from two-tailed Mann-Whitney U test. **(h)** Ki67 staining of hair matrix. **(i)** Ex vivo live imaging of human HF (14.5h) identifying cell division events (green dots) in the matrix in a similar distribution to Ki67 staining. Red dotted lines demarcate IRS, cortex and medulla layers. **(j-k)** Live imaging tracking of cell movements for basal and differentiated matrix keratinocytes showing minimal movement in the COL17A1+ basal populations followed by rapid upward cell flow in differentiated HF layers. Red dotted lines demarcate IRS, cortex and medulla lineages. **(l)** Tracked cell displacement over 20.5 hours for matrix basal progenitors (lower and upper COL17A1, medulla) revealing minimal movement in the lower and upper COL17A1 populations. ***denotes statistical significance from two-tailed Mann-Whitney U test.

To define matrical lineage organization *in situ*, we performed spatial mapping of the high-resolution single-cell subclusters onto ST datasets. This revealed a highly compact, layered architecture in the lower HF bulb, consistent with histologic organization (Fig. 5c). The superimposed spatial transcriptome allowed direct visualization of gene expression patterns and lineage relationship in place (Fig. 5c and Extended Data Fig. 7b). The outer perimeter of the lower HF was flanked by LPC cells, positioned at the distal bulb, abutting the lower DP, and extending toward the turning point of the hair bulb, reminiscent of the murine organization^69^. Along the matrix-DP border, LPC cells were followed by matrix basal cells that extended upward until the medulla population at the DP apex (Fig. 5c). Although *COL17A1* expression levels were comparable between LPC and matrix basal progenitors at both mRNA and protein levels (Fig. 5b and Extended Data Fig. 7a), matrix basal cells further mapped to three spatially segregated and transcriptomically distinct populations (Supplementary Table 9). We termed these lower COL17A1+ basal progenitors, upper COL17A1+ basal progenitors, and medulla progenitors, based on their position along the matrix-DP border (Fig. 5d). Spatially, the lower COL17A1+ cluster mapped to the basal region below the line of Auber, a transverse landmark across the anagen bulb at the widest level of the DP^70^, and above the LPC zone. The upper COL17A1+ cluster mapped above the line of Auber and intermingled with melanocytes responsible for HS pigmentation. Medulla progenitors occupied the basal layer between the upper COL17A1+ zone and the DP apex (Fig. 5d).

More differentiated IRS and cortex lineages extended away from their corresponding basal compartments and largely followed a radially concentric arrangement. The early IRS population dovetailed with the lower COL17A1+ basal zone and subsequently branched into three distinct IRS layers along the HF axis (Fig. 5c and Extended Data Fig. 7b). By contrast, the HS cuticle and cortex emerged from the upper COL17A1+ basal zone and progressed upward with increasing maturation (Fig. 5c and Extended Data Fig. 7b). Interestingly, *FOXQ1+* medulla layers showed a distinct pattern: they were largely confined to the basal layer along the basement membrane and only delaminated into multi-layered structures near the DP apex (Fig. 5c and Extended Data Fig. 7b-c).

This organization, where multiple lineage trajectories arise from discrete COL17A1+ basal compartments, differs from conventional models in which a broad Ki67+ proliferative zone spans the entire cup-like matrix surrounding the DP up to the line of Auber, with differentiation initiating upon cell cycle exit^70–72^. Indeed, without regressing cell cycle genes, scRNA-seq analyses recovered a prominent proliferative matrix cluster (separable into proliferative IRS and cuticle/cortex subsets) marked by *MKI67, TOP2A,* and related cell cycle transcripts, which formed distinct cell clusters in the UMAP (Extended Data Fig. 7d-e). After cell cycle gene regression (Methods), these proliferative cells redistributed across early differentiation clusters within their respective (IRS and cortex) lineage branches in the single-cell UMAP (Extended Data Fig. 7f-g). Consistent with this lineage-specific assignment, spatial mapping of these highly proliferative cells localized them to the regions of hair club that correspond to the IRS and cortex lineages, respectively (Extended Data Fig. 7h). These results suggest that proliferation status, rather than lineage relationship, dominated single-cell clustering without the proper correction (Extended Data Fig. 7d), and that proliferative matrix cells already exhibit lineage-biased transcriptional programs beyond cell cycle genes (Extended Data Fig. 7f-h).

In spatial mapping, proliferative cells were primarily enriched within early differentiation clusters rather than within COL17A1+ basal populations (Extended Data Fig. 7h-i), suggesting that they function as lineage-committed transient amplifying cells (TACs). To compare these highly proliferative TACs with their corresponding basal progenitors, we computed a per-cell adhesion module score based on statistically significant ECM ligand-receptor interactions between matrical populations and the DP, identified by CellChat^73^ (Methods). As expected, LPC cells and both COL17A1+ basal populations showed the highest adhesion scores. Medulla progenitors, however, displayed significantly lower adhesion scores (Fig. 5f) despite high COL17A1 expression at both mRNA and protein levels (Fig. 5e and Extended Data Fig. 7a). Early differentiation populations overlapping with TACs displayed intermediate adhesion scores, whereas terminally differentiated cells showed the lowest scores (Fig. 5f). We next examined proliferation across the three basal compartments. Lower and upper COL17A1+ basal populations showed high proliferation scores, whereas medulla progenitors showed markedly reduced proliferation score (Fig. 5g), a pattern corroborated by Ki67 mRNA and protein staining (Fig. 5h and Extended Data Fig. 7i). Notably, within both lower and upper COL17A1+ basal populations, adhesion and proliferation were inversely correlated (Extended Data Fig. 7j), consistent with a continuum in which strongly adherent basal cells are relatively less proliferative, whereas less adherent progeny transition into a proliferative TAC-like state.

To validate these spatial inferences and directly observe lineage behavior, we performed time-lapse live imaging (15-20.5 hours) of the matrix region in freshly isolated human scalp hair^74^ (Movie S1) and tracked single-cell dynamics and lineage relationship *in situ* (Fig. 5i-k). Consistent with our *in situ* detection of *Ki67* mRNA (Visium HD data, Extended Data Fig. 7i), single cell mapping (Extended Data Fig. 7h) and KI67 IF results (Fig. 5h), delaminated IRS and cortex progenitors were highly proliferative, whereas delaminated medulla progenitors did not divide (Fig. 5i). Strikingly, basal cells in the lower and upper COL17A1+ zones were largely immobile, with an average displacement of 5.4 um over 20 hours, based on single-cell tracking, whereas basal cells within the medulla progenitor zone migrated upward along the basement membrane with an average displacement of 17.9 um over the same period (Fig. 5j-l). Furthermore, basal cells in the lower and upper COL17A1+ zones occasionally delaminated or divided to generate progenitors that embarked on IRS and cortex differentiation trajectories, respectively (Movie S1). These progenitors displayed elevated cell division activities, compared with their basal cell counterparts (Fig. 5i). In striking contrast, dividing cells were rarely observed among medulla basal progenitors. Instead, these progenitors generally moved upward and delaminated near the DP apex to form a corn-shaped medulla differentiation zone (Fig. 5j-k) and Movie S1). Interestingly, tracking individual IRS and cortex progenitors immediately after delaminating revealed layered but limited cell movement patterns (Fig. 5j-k). Upon reaching the upper matrix, their upward movement accelerated and reached the speed of ∼6.8um/h. Remarkably, the layered cell lineage organization was evident across all three major matrix lineages - IRS, cortex and medulla – throughout the imaging interval (Fig. 5j-k and Movie S1), mirroring the spatial mapping results.

Together, these spatially resolved analyses integrate high-resolution mapping and live imaging to support a unifying organizational principle for the human hair bulb. Along the matrix-DP interface, COL17A1+ basal progenitors and their delaminating progenitors co-reside but differ in adhesion and proliferative states. Progenitors with the strongest adhesion programs remain anchored to the basement membrane and divide less frequently, consistent with a more stable stem/progenitor reservoir. As adhesion programs attenuate, cells divide or delaminate, enter a rapid transit amplification state, and commit to IRS or HS trajectories that fuel fast yet orderly hair growth. In contrast, medulla layers arise from a layer of less proliferative basal cells that migrate along the basement membrane before delaminating near the DP apex to initiate terminal differentiation ((Extended Data Fig. 7k).

### Analysis of Primed Chromatin-Gene Expression Relationship In Situ

Although lower and upper COL17A1+ basal populations shared core adhesion and proliferation programs, they were already transcriptionally distinct (Supplementary Table 9). Notably, each population expressed differentiated lineage markers despite their basal position at the matrix-DP interface: lower COL17A1+ cells expressed IRS genes including *GATA3*, *SHH* and *KRT28*, whereas upper COL17A1+ cells expressed HS-associated genes such as *KRT35*, *KRT85* and *CPA6* (Fig. 6a). Consistent with early fate commitment, spatial expression patterns further revealed rapid induction of lineage-specific transcripts in delaminated early progenitors, such as *KRT32* (Extended Data Fig. 8a–b). These observations led us to hypothesize that matrix basal progenitors are lineage-primed, with fate bias encoded in chromatin accessibility before the transcription of differentiation genes.

**Fig. 6:**
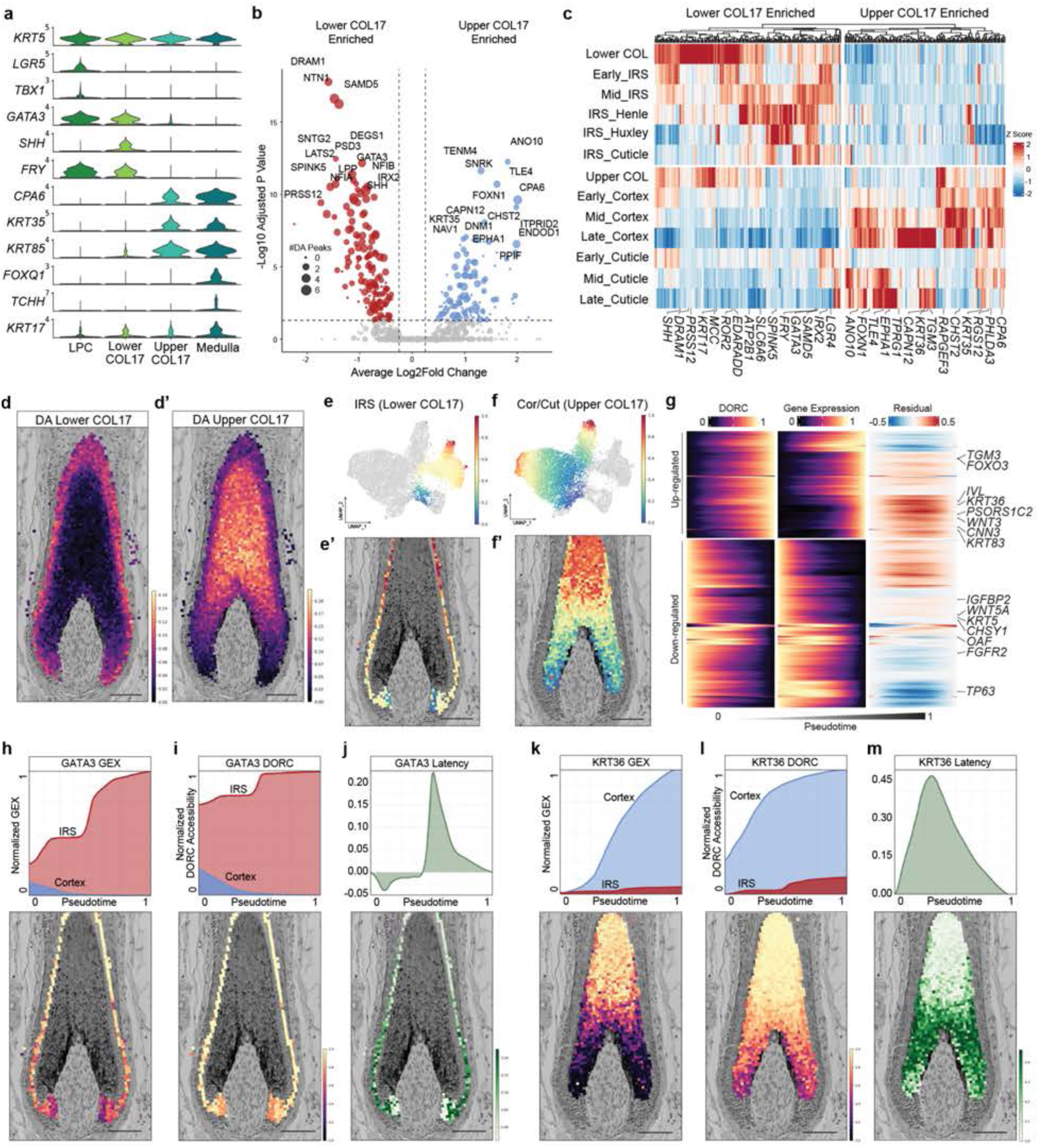
Multimodal Analysis of Chromatin and Gene Expression Dynamics in Human Matrix *in situ*. **(a)** Violin plot of LPC (*KRT5, LGR5, TBX1*), IRS (*GATA3, SHH, FRY*), cortex (*CPA6, KRT35, KRT85*), and medulla (*FOXQ1, TCHH, KRT17*) lineage marker gene expression for *COL17A1*+ basal populations. **(b)** Volcano plot of genes with differentially accessible (DA) domains of regulatory chromatin (DORC) between lower and upper COL17A1+ matrix basal progenitors. **(c)** Hierarchical clustering heatmap of pseudobulked gene expression for differentially accessible DORC genes between lower and upper COL17A1+ matrix basal progenitors. **(d-d’)** Spatial visualization of gene module activities with H&E overlay of the lower hair follicle for lower COL17A1- (d) and upper COL17A1-enriched (d’) DA DORC genes. Scale bar: 100µm. **(e-e’)** Palantir pseudotime heatmap of the lower COL17A1-IRS trajectory (e) and visualization of the single-cell pseudotime values mapped onto ST data with H&E overlay of the hair matrix. Scale bar: 100µm. **(f-f’)** Palantir pseudotime heatmap of the upper COL17A1-cortex/cuticle trajectory (f) and visualization of the single-cell pseudotime values mapped onto ST data with H&E overlay of the hair matrix. Scale bar: 100µm. **(g)** Hierarchical clustering of smoothed and normalized DORC accessibility (left), RNA expression (middle), and DORC-RNA residuals (right) across the cortex trajectory for upregulated- and downregulated- expressed genes. **(h-j)** Line plot of *GATA3* gene expression (GEX) (h) and chromatin accessibility (DORC) (i) for the IRS and cortex/cuticle lineages as well as the difference between chromatin accessibly and gene expression, defined as gene latency (j). Below each panel is a visualization of the single-cell mapped corresponding values onto ST data with H&E overlay of the hair matrix. Scale bar: 100µm. **(k-m)** Line plot of *KRT36* gene expression (GEX) (k) and chromatin accessibility (DORC) (l) for the cortex/cuticle and IRS lineages as well as the difference between chromatin accessibly and gene expression (latency) (m). Below each panel is a visualization of the single-cell mapped corresponding values onto ST data with H&E overlay of the hair matrix. Scale bar: 100µm.

To test whether fate bias is established at the chromatin level, we leveraged co-profiled single-cell multiome data enriched for hair matrix compartments and applied an open chromatin peak-gene linkage framework to define domains of regulatory chromatin (DORCs), gene-centric collections of cis-regulatory elements whose aggregate accessibility predicts associated transcription^75^. This analysis identified 1,485 highly regulated genes with robust regulatory support (≥8 linked peaks), enabling quantitative comparisons of locus-level accessibility between progenitor compartments. Differential DORC accessibility analysis between lower and upper COL17A1+ progenitors identified 286 genes with lineage-biased regulatory landscapes with 154 enriched in the lower compartment and 132 enriched in the upper compartment (Fig. 6b and Supplementary Table 10). Known lineage-associated loci exhibited the expected polarization, with IRS regulators and markers (*GATA3*, *SHH*, *NFIA/B*, *SAMD5*) showing higher DORC accessibility in lower COL17A1+ cells, and HS-associated genes (*CPA6*, *FOXN1*, *KRT35*, *KRT36*) enriched in upper COL17A1+ cells (Fig. 6b). Furthermore, newly identified genes that were linked to differentially accessible DORCs also segregated transcriptionally along IRS vs HS differentiation trajectories - pseudobulk expression of these loci robustly separated IRS and HS lineages, consistent with their compartment-specific chromatin states (Fig. 6c).

Several loci exhibited a canonical priming pattern in which DORC accessibility was already elevated in basal progenitors, whereas transcript accumulation occurred primarily in more differentiated states (for example KRT36, activated later in cortex/cuticle). To visualize this regulatory priming directly in tissue space, we built gene modules from lower- and upper-enriched DORC loci and projected module scores onto the spatial transcriptomics maps. This revealed a striking spatial separation of lower- vs upper-primed chromatin programs that aligned with their respective IRS and HS domains (Fig. 6d–d’), indicating that fate-restricted regulatory architectures are spatially organized and detectable *in situ*, reminiscent of transcriptional divergence.

Consistent with this model, chromatin motif analysis (chromVAR) identified distinct motif enrichments between the two basal compartments, with lower progenitors enriched for NFI and GATA family motifs and upper progenitors enriched for Homeobox family motifs (Extended Data Fig. 8c). At the protein level, co-IF staining further supported polarized lineage regulation, with GATA3 preferentially marking IRS layers and HOXC13 marking HS cortex/cuticle in mutually exclusive matrical domains extending to the basement membrane (Extended Data Fig. 8d). Notably, this spatial partitioning of regulatory chromatin dynamics and lineage-associated transcription was concordant with the lineage trajectories visualized by live imaging (Fig. 5j-k).

We next asked how chromatin priming relates to the unusually rapid differentiation dynamics of human matrix lineages. Using Palantir^76^, we reconstructed pseudotime trajectories for IRS and HS cortex/cuticle lineages, rooting each trajectory in the lower or upper COL17A1+ basal progenitor population, respectively. Projecting pseudotime onto spatial coordinates revealed accelerated differentiation shortly after delamination from the basement membrane, with trajectories extending from basal origins to mature states several hundred microns (∼30–40 cells) away (Fig. 6e-e’ and 6f-f’). To integrate chromatin and transcription dynamics, we smoothed gene expression and DORC accessibility over pseudotime and evaluated concordance and temporal offsets between open chromatin accessibility and RNA accumulation. Across the cortex/cuticle trajectory, many late differentiation genes, including those from keratinization-associated loci, exhibited accessibility increases preceding transcriptional upregulation, consistent with enhancer opening before RNA induction (Fig. 6g). Conversely, some basal-state genes (for example TP63) showed rapid loss of enhancer accessibility shortly after delamination followed by decreases in RNA, consistent with chromatin-level shutoff at the onset of differentiation (Fig. 6g).

Detailed analyses of three loci further illustrated these principles. In the IRS trajectory, *GATA3* showed elevated baseline accessibility and detectable basal expression in lower COL17A1+ progenitors, followed by additional accessibility gains accompanying progressive RNA and protein induction specifically along IRS differentiation (Fig. 6h-j and Extended Data Fig. 8d-e). In the HS trajectory, *KRT36* exhibited elevated baseline accessibility in upper progenitors despite minimal RNA expression at the trajectory root, followed by rapid transcriptional activation and a gradual increases in accessibility during cortex/cuticle maturation (Fig. 6k-m and Extended Data Fig. 8f). Conversely, genes comprising the matrical basal transcriptional program were rapidly downregulated along both the lower and upper COL17A1+ lineages upon differentiation. For example, TP63 was highly expressed at the root in both trajectories (Extended Data Fig. 8g-h) and exhibited elevated DORC accessibility. Shortly after delamination, TP63 DORC accessibility decreased sharply, followed by a lagged decline in mRNA abundance across both lineages (Extended Data Fig. 8i-j), yielding a negative latency (Extended Data Fig. 8k).

We noted that these predicted expression profiles derived from spatial mapping of single-cell transcriptome were highly concordant with the native Visium HD signals *in situ* (Extended Data Fig. 8e-g), supporting the accuracy of the integration framework. Finally, direct inspection of DORC-linked peaks across pseudotime-binned cells recapitulated these locus-level dynamics, including early accessibility of *KRT36*-linked regulatory peaks in upper but not lower progenitors and progressive accessibility increases during HS differentiation, as well as late IRS cuticle gene priming exemplified by *KRT73* (Extended Data Fig. 8l–m).

Together, these *in situ* multimodal analyses demonstrate that seemingly similar COL17A1+ basal cells are epigenetically partitioned into IRS- and HS-primed progenitor populations, respectively. Their fate-biased chromatin landscapes facilitate rapid lineage-specific transcription and spatially organized differentiation, providing a mechanistic framework for how the human matrix sustains continuous hair production through pre-patterned regulatory programs.

### Atlas-Guided Analysis of Inflammatory Hair Loss Diseases

Having established a spatially resolved reference atlas of healthy human scalp, we next evaluated its utility for accurately assessing disease-associated changes in cell composition and state with the goal of improving spatial interpretation of patient scRNA-seq profiles and nominating pathways relevant to diagnosis and treatment. To this end, we generated scRNA-seq datasets from four lesional punch biopsies collected from treatment-naïve, early-to-mid stage lichen planopilaris (LPP) patients in our hair loss clinics. As a comparison, we analyzed four alopecia areata (AA) datasets with matched controls from a prior publication^77^.

LPP and AA represent prototypical inflammatory scarring and non-scarring alopecias, respectively^78,79^. Using the reference atlas as a scaffold, we constructed a shared scVI model followed by scANVI-based integration and label transfer to harmonize cell identities across healthy and disease samples^80^ (Fig. 7a). Aggregated cell-type proportions revealed expansion of lymphoid and myeloid compartments in both AA and LPP relative to controls (Fig. 7b), consistent with immune-cell infiltration as a core feature of inflammatory alopecias.

**Fig. 7:**
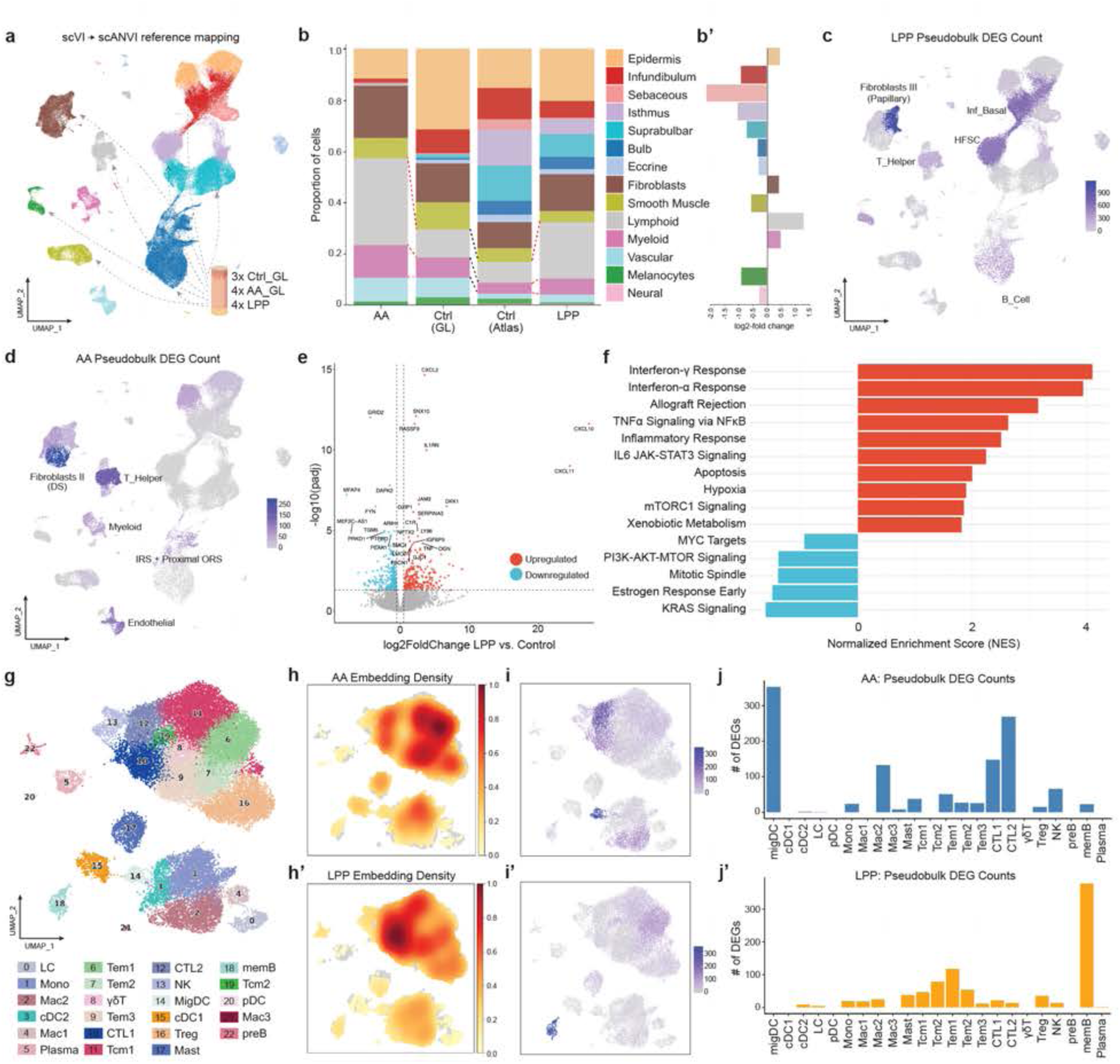
Atlas Integration and Reference Mapping of Inflammatory Hair Loss. **(a)** Reference mapping of published control (Ctrl_GL), alopecia areata (AA_GL) and new lichen planopilaris (LPP) scRNA-seq datasets. **(b)** Stacked bar plot showing cell composition by sample source and disease condition after atlas integration. **(b’)** Cell type composition changes in LPP samples compared to control showing reduction of upper HF epithelial compartments and enrichment of immune populations. **(c-d)** Feature plots of pseudobulk differentially expressed gene (DEG) counts for LPP (c) and AA (d) revealing the key affected cell populations in each disease state. **(e)** Volcano plot of DEGs in HF-SCs in LPP compared to control. **(f)** Gene set enrichment analysis of LPP HF-SC DEGs reveals elevated interferon response among other pathways. **(g)** UMAP representation of 23 immune cell subtypes in healthy, AA, and LPP scalp tissue. **(h-h’)** Cell embedding density of AA and LPP samples. **(i-i’)** UMAP visualization of pseudobulk DEG counts for AA and LPP immune cells compared to control. **(j-j’)** Bar plot of pseudobulk DEG counts for AA and LPP by identified immune subsets.

Because sample-to-sample variation can confound compositional comparisons, we next applied a Bayesian model to test differential cell abundance in LPP relative to healthy^81^, both of which were generated using the same optimized workflow. This analysis confirmed the increased lymphoid and myeloid populations and, importantly, revealed a significant depletion of upper HF epithelial compartments, most prominently the sebaceous lineage and the HF-SC region (Fig. 7b’), while lower HF compartments (suprabulbar and bulb) were comparatively preserved, consistent with biopsy sites selected from clinically early-to-mid disease with relatively maintained hair density (Fig. 7b’). These atlas-resolved compositional changes mirror classic histopathology in LPP, including sebaceous gland atrophy and inflammatory targeting of the upper follicle/HF-SC region that ultimately compromises regenerative capacity and promoting scarring hair loss^82^.

We next leveraged our integrated cell type annotations between control and diseased samples to identify cell compartments that were most strongly affected in AA and LPP, respectively. Pseudobulked DEG analysis revealed that upper HF epithelial compartments, a subcluster of fibroblasts and B cells were most affected in LPP whereas a distinct subset of fibroblasts, T helper cells and vascular cells were most affected in AA (Fig. 7c-d). Because both diseases showed perturbations in fibroblasts, this indicated stromal participation or response to the underlying pathology in these distinct disorders. Strikingly, guided by our spatially resolved fibroblast mapping (Fig. 3a-c), the Fibroblasts III population composed of papillary and upper perifollicular fibroblasts were identified as the most affected in LPP (Fig. 7c), whereas the Fibroblasts II population that contains two dermal sheath subsets was identified as the most perturbed in AA (Fig. 7d). These results are consistent with canonical locations of inflammation on histology^82^ and highlight the ability to glean spatially relevant information when pairing our atlas with dissociation-based scRNAseq data.

Although LPP is a prototype scarring alopecia and often considered a disease characterized by HF-SC dysregulation and loss of HF immune priviledge^83^, compartment-resolved transcriptional consequences within human HF-SCs remain incompletely defined. Leveraging our spatially integrated healthy and diseased annotation, we next determined the gene expression changes in HF-SCs in LPP. Compared to control, LPP HF-SCs upregulated interferon-inducible chemokine, most prominently CXCL10/11, which are high-affinity ligands of CXCR3 that recruit activated T cells and amplify type I inflammatory circuits^84^ (Fig. 7e). Gene set enrichment analysis (GSEA) further uncovered marked induction of type I/II interferon response programs and antigen-presentation/immune-activation modules, including B2M and MHC I components, consistent with immune-privilege collapse (Fig. 7f; Allograft Rejection). In parallel, pro-apoptotic and cell death-associated pathways, including FAS, BAX, and various caspases, were significantly upregulated (Fig. 7f), aligning with the observed depletion of the HF-SC compartment and with prior evidence for inflammation-associated HF-SC injury in LPP (Fig. 7b).

Although both AA and LPP are inflammatory alopecia, the precise immune milieu of each condition remains poorly defined. We next investigated the utility of our healthy scalp atlas as the framework to pinpoint changes in the immune compartment in each disease. Using the healthy control scalps as reference, we integrated AA and LPP immune subsets and identified 23 unique immune cell sub-populations. Using a combination of canonical immune marker genes and logistic regression classifiers (see Methods) (Extended Data Fig. 9 and Supplementary Table 11), we determined nuanced identities to these immune clusters including multiple subsets of cytotoxic and helper T cells, macrophages, dendritic cells, and B cells (Fig. 7g). Notably, the cell embedding density of AA and LPP samples suggested a preponderance of T effector memory cells in AA and increased cytotoxic T cells in LPP, respectively (Fig. 7h-h’). When examining differentially expressed genes, however, migratory dendritic cells (migDC) and cytotoxic T cells (CTLs) were most perturbed in AA (Fig. 7i-j), reflecting the typically brisk inflammation leading to HF matrix destruction (REF). In sharp contrast, memory B cells were the most perturbed in their DEGs in LPP whereas gene expression changes in migDCs and CTLs exhibited few changes (Fig. 7i’-j’). The DEG list of these memory B cells was enriched for genes involved in canonical B cell activation (TNF, TRAF1, IRF4, MIR155HG, CD44)^85^, NF-kB signaling (BIRC3, NFKBIA/NFKBIZ)^86^, cell survival (MCL1, BCL2A1)^87^, and tissue retention (ITGA4)^88^ (Supplementary Table 12). While T cell-driven immune privilege collapse is well appreciated in LPP, these data nominate an additional, disease-associated B cell activation axis that has been comparatively under-characterized in scarring alopecia and may contribute to chronicity and HF-SC attrition. Taken together, the reference atlas-guided integration delineates divergent, spatially interpretable inflammatory programs in AA vs LPP across stromal, epithelial and immune compartments, providing a molecular and cellular foundation for mechanistic studies and the rational stratification of therapeutic targets.

## Discussion

Here, we constructed a high-fidelity, spatially resolved reference atlas of healthy human scalp by integrating scRNA-seq, chromatin accessibility and deep-coverage spatial transcriptomics. By anchoring single-cell transcriptome to *in situ* coordinates through high-dimensional ST data, the atlas resolves epithelial, mesenchymal and immune states across the entire hair unit, including populations that have been difficult to capture robustly in prior human datasets. Beyond serving as a community resource, this framework links spatial position to regulatory state and lineage progression, enabling mechanistic interpretation of patterned human tissue architecture in health and disease.

Our result indicates that transcriptome depth per spatial unit is a key determinant of mapping fidelity in complex and dense tissues such as hair matrix. Although many spatial platforms emphasize miniaturization^89^, insufficient gene capture limits robust deconvolution and integration^8^. In our data, modest reductions in RNA integrity lowered gene capture and measurably degraded spatial mapping performance. Reducing the genome wide gene capture to the 5k panel of Xenium also compromised the mapping fidelity, underscoring the importance of high gene dimensionality as a critical determinant for accurate spatial assignment. Moreover, current cell segmentation methods, especially in high cell density tissues, are imperfect, and erroneous cell segmentation and transcript assignments have been shown to confound downstream analyses^90^.

Biologically, the atlas delineates a spatially restricted human HF-SC niche at the APM-HF junction and provides a comprehensive marker repertoire for improved isolation and functional studies. Spatial mapping resolves stromal heterogeneity into anatomically distinct fibroblast states, including papillary, reticular, dermal sheath and perineural populations, thereby enabling interpretable hypotheses about microenvironmental functions and cell-cell communication. In parallel, we resolve compartment-specific melanocyte organization, including two matrical states with divergent melanogenic programs consistent with the high pigment demand of continuous hair growth.

Most notably, integrating deep spatial mapping with multiome profiling and live imaging reveals an organizational principle of the human hair matrix. Spatial reconstruction identifies segmented COL17A1+ basal progenitor domains along the matrix-DP interface that seed lineage-specific differentiation trajectories. Proliferation is enriched in delaminated, early differentiation states, consistent with lineage-committed transient amplifying cells, while chromatin accessibility in basal progenitors is already fate-biased, indicating epigenetic priming before transcriptional divergence. Live imaging, uniquely feasible in human follicles, provides orthogonal validation by directly visualizing compartment-specific cell behaviors and lineage dynamics that mirror spatially inferred trajectories. Such architecture - chromatin-poised stem/progenitor cells anchored to an adhesive niche, TACs as an expandable intermediate, and spatially stratified differentiation - may represent a common feature in human tissues that couples rapid proliferation with robust differentiation while maintaining long-term structural integrity.

Finally, using the atlas as a reference coordinate system, we map AA and LPP scRNA-seq profiles into defined spatial and lineage contexts, revealing disease-specific perturbations across fibroblast niches, epithelial compartments and immune programs. Together, this work establishes both a high-resolution reference for human scalp biology and a generalizable strategy for connecting transcriptomically defined cell state, tissue architecture and cell behavior in health and disease.

## Methods

### Human Scalp Tissue Acquisition and Consent

Healthy human scalp tissue samples were obtained either from discarded scalp tissue from patients undergoing routine dermatologic surgeries or from 4-6 mm punch biopsies from healthy volunteers. Scalp from the surgical site and donor biopsy sites were examined by a board-certified dermatologist to verify the absence of clinically-evident disease states prior to enrollment in the study. Samples were collected with informed consent within the Northwestern University Institutional Review Board approved protocol. Immediately after excision/biopsy, the scalp samples were placed in MACS Tissue Storage Buffer (Miltenyi Biotech) at 4°C on ice until downstream processing. Samples were stored for no longer than 4 hours prior to single cell dissociation.

### Single-Cell Dissociation

Samples for single cell dissociation were rinsed in ice-cold 1x HBSS prior to processing with either 4 mm punch biopsy for large discarded surgical tissues, hair follicular unit extraction (FUE) using a 0.8 mm motorized rotary punch tool, or micro-dissected for lower hair follicle isolation by orienting the tissue with the dermal side-up. The resulting tissue was then diced in a grid-like fashion into submillimeter pieces with a scalpel blade on ice. The diced tissue was then dissociated using the Whole Skin Dissociation Kit (Miltenyi Biotec) per manufacturer protocol. In brief, samples were incubated in 0.5 ml of dissociation buffer containing described volume of enzyme P, D, and A for 3 hours at 37°C under gentle agitation. Thereafter, 0.5 ml of RPMI with 10% FBS was added to each sample and then mechanically dissociated using the GentleMACS Tissue Dissociator with the h_skin_01 program. Following enzymatic and mechanical tissue dissociation, samples were gently triturated with 10 ml of RPMI with 10% FBS and filtered through a 40 µm cell strainer. The filtrate was centrifuged for 10 minutes at 300g at 4°C to pellet the primarily stromal cell fraction. Concurrently, the undigested tissue debris captured in the cell strainer were washed out using 3 ml of pre-warmed 0.05% trypsin-EDTA and incubated for 12 minutes at 37°C under gentle agitation to further dissociate the keratinocyte-rich fraction of the tissue, particularly the hair follicle associated structures. The trypsinized sample was then neutralized with 10 ml of RPMI with 10% FBS and filtered through a 40 µm cell strainer. The cell suspension was centrifuged for 10 minutes at 300g at 4°C to pellet the cells. After removing the supernatant, both pellets were then resuspended in PBS with 1% BSA and combined to further wash the cells. The combined cell suspension was then centrifuged and resuspended in the appropriate volume of PBS with 1% BSA for the single cell RNA-sequencing pipeline or nuclei isolation. All samples were stained with trypan blue for viability assessment and counted using both a Countess II automated hemocytometer (Invitrogen) as well as a manual hemocytometer as cross-validation. All cells used for experiments related to the current study were freshly isolated without undergoing freeze-thaw.

### Single-Nuclei Isolation

After tissue dissociation and single cell isolation, samples to be used for single-cell multiomics (combined scRNA-seq and scATAC-seq) studies were processed for nuclei isolation following the Nuclei Isolation for Single Cell ATAC Sequencing protocol (10x Genomics) with the exception of the addition of Protector RNase Inhibitor (Roche) to all lysis and wash buffers. In brief, approximately 500,000 cells were pelleted and resuspended in cell lysis buffer for 5 minutes. The resulting nuclei suspension was centrifuged for 5 minutes at 500g at 4°C to pellet the nuclei and washed twice with nuclei wash buffer followed by centrifugation and pellet resuspension. The final nuclei suspension was filtered through a 40 µm Flowmi cell strainer (Sigma Aldrich) prior to final centrifugation and pellet resuspension in dilute nuclei buffer for automated and manual nuclei counting and direct visualization of nuclei quality.

### Single-Cell RNA-Sequencing

Following single cell isolation, cell count determination, and live/dead ratio assessment, samples with greater than 85% live cells were used for subsequent single-cell RNA-sequencing. The required volume of cells was loaded onto the Chromium controller (10x Genomics) for single cell encapsulation, targeting 10,000 cells per sample for Next GEM single cell 3’ RNA v3.1 and 20,000 cells per sample for GEM-X 3’ RNA v4. Sequencing libraries were constructed according to 10x Genomics protocol with unique Illumina indices. The resulting single cell RNA libraries were sequenced on the Illumina NovaSeq X Plus platform with an average of 45-50K read-pairs per cell. BCL-convert was used to generate fastq files from raw sequencing reads. The resulting files were mapped with the 10x Genomics CellRanger pipeline (version 8.0.1) to human reference genome GRCh38 (v2024-A) using the CellRanger count function.

### Multiomics Sequencing

Following nuclei isolation and quantification, the required volume of nuclei targeting 10,000 nuclei were loaded onto the Chromium controller using Chip J for nuclei encapsulation. Nuclear chromatin transposition (for ATAC) and cDNA amplification (for GEX) were performed according to manufacturer instructions (10x Genomics). The resulting scRNA and scATAC libraries were sequenced on the Illumina NovaSeq X Plus platforms targeting an average of 45-50K read-pairs per cell for GEX and 60K read-pairs per cell for ATAC. BCL-convert was used to generate fastq files from raw sequencing reads. The resulting files were mapped with the 10x Genomics CellRanger-ARC pipeline (version 2.0.2) to human reference genome GRCh38 (v2024-A) using the CellRanger-ARC count function.

### Spatial Transcriptomics Sample Preparation and Sequencing

Patient samples used for spatial transcriptomics were immediately placed in 4% PFA for tissue fixation at 4°C for 6-12 hours depending on the thickness of the tissue (∼2 hours per mm). After fixation, the tissue was trimmed to remove excess exposed hair shafts protruding above the epidermis and subcutaneous adipose tissue. The tissue samples were then washed with PBS and acclimated in 50% ethanol for 30 minutes prior to transitioning to 70% ethanol for storage at 4°C. For paraffin processing and embedding, tissue samples were placed in standard histology cassettes and dehydrated and wax-embedded using a Leica HistoCore processor. The resulting paraffin infiltrated tissues were then embedded in paraffin using a FFPE tissue embedding center (Leica). Samples for Visium HD were sectioned at 5 µm thickness and floated in a 42°C water bath containing RNase-free water for 1 minute prior to adhesion to Nexterion 3D hydrogel-coated slides (Schott Inc). Subsequent steps including deparaffinization, H&E staining, destaining, decrosslinking, probe hybridization, and CytAssist transfer to Visium HD slides were performed according to manufacturer instructions. Four 5 µm sections from the start and end of tissue sectioning were collected for RNA isolation using the RNeasy FFPE kit (Qiagen) per manufacturer protocols. The isolated RNA was analyzed for RNA quality by the DV200 metric using the RNA 6000 Pico kit (Agilent) on a Bioanalyzer 2100 instrument (Agilent). A minimum DV200 of 40% was required prior to proceeding with the Visium HD workflow.

Post-CytAssist probe amplification and Illumina sequencing library construction were performed according to the 10x Genomics published protocols. Resulting spatial transcriptomics libraries were sequenced on the Illumina NovaSeq X Plus platform with a target of 1,000M read-pairs/capture area, with adjustments made to account for the amount of tissue coverage. BCL-convert was used to generate fastq files from raw sequencing reads. The resulting files were mapped with the 10x Genomics SpaceRanger pipeline (version 3.1.2) to human reference genome GRCh38 (v2024-A) using the SpaceRanger count function.

Efforts were made to ensure all the steps in processing remained as RNase-free as possible, with utilization of RNase-free reagents and labware when possible and extensive cleaning of surfaces with RNaseZap (Invitrogen) to minimize RNA degradation during processing. FFPE blocks were stored at room temperature in a dehydrator to minimize RNA degradation.

### Processing and Quality Control of scRNA-seq Datasets

Removal of ambient RNA and detection of non-empty droplets were performed using CellBender (v.0.3.0)^91^ using the CellRanger raw_feature_bc_matrix.h5 file as input. Each sample was processed using a learning rate of 0.00005 with all other parameters set to default values. CellBender-filtered gene expression matrices were then imported into Seurat (v5.1.0)^92^ for further downstream analyses. A two-step strategy was employed to remove low-quality cells. First, a two-factor linear mixture model (miQC v1.12.0)^93^ was used to model the relationship between the number of detected genes per cell and the mitochondrial UMI percentage. Cells with a posterior probability >0.75 for the inferred ruptured population were considered dead/dying and excluded from downstream analysis. Next, outlier cells with <800 or >8,000 detected genes were removed from downstream analysis. Finally, scDblFinder (v1.18.0)^94^ was used on each sample using the default parameters to remove doublets prior to integration.

### Preprocessing and Quality Control of Multiomics datasets

Ambient RNA removal and droplet calling was performed for the scRNA-seq modality of multiomics datasets as described above. scATAC-level thresholding for cells passing RNA-level filtering was performed with ArchR (v1.0.2)^95^. Nuclei containing fewer than 1,000 fragments, a TSS enrichment score <4, nucleosome ratio <2, or more than 2% of reads falling into blacklisted regions were removed. Finally, doublet identification and removal were performed for each sample using the paired scRNA data with scDblFinder as described above.

### Integration of Multiomics and scRNA-seq Datasets

Following sample preprocessing and QC, all scRNA-seq and multiomics samples were merged into a single Seurat v5 object for downstream analyses. Counts were normalized to a library size of 10,000 and log-transformed using the NormalizeData() function. Cell cycle scoring was performed using the log-normalized count matrix using the CellCycleScoring() function. The top 2,000 highly variable genes (HVGs) were computed across all cells with the FindVariableFeatures() function using the “vst” method. Integration was performed using an scVI model (n_layers=2, n_latent=50, gene_likelihood=”nb”) implemented through scVI-tools (v1.3.0)^96^ with individual samples as the batch key and the following categorical covariates (Donor ID, platform version, and sex) and continuous covariates (feature count, UMI count, mitochondrial gene percentage, and ribosomal gene percentage). To mitigate artifacts in gene expression associated with sample processing, a panel of published single-cell dissociation response genes^97^ was removed from consideration for integration and clustering. The log-normalized expression values of each cell cycle and dissociation response gene were included as additional continuous covariates to mitigate their effects on clustering. Training was performed for a maximum of 500 epochs or until the model reached convergence. Dimensionality reduction and Louvain clustering were performed using the scVI latent space using the FindNeighbors(), FindClusters() and RunUMAP() functions in Seurat. Cluster markers were initially identified using the FindAllMarkers() function for general cluster annotation. Subsequent analyses of individual higher-resolution subsets were performed first by repeating the highly variable feature selection, scVI integration, and downstream dimensionality reduction, and clustering as described above.

### Atlas Level Integration and Label Transfer Using scANVI

Additional scRNA-seq datasets (control and disease samples) were processed using the previously described computational pipeline for ambient RNA, low-quality cell, and heterotypic doublet removal. The processed datasets were then concatenated with the annotated atlas with their annotations as “Unknown” to create a combined object. A new scVI model was calculated using the previously described parameters to create a joint embedding that incorporated the new query cells. Finally, an scANVI model was calculated from the joint scVI model to transfer the cell type annotations from the atlas to the query datasets.

### Analysis of Multiomics Paired scATAC-seq Chromatin Accessibility Datasets

High-resolution cell type annotations for single-cell multiomics samples were generated as described above. To maximize granularity in peak-calling across cell types, nuclei were first grouped by high-resolution subclusters prior to peak-calling. Resulting chromatin accessibility profiles were pseudobulked by cell type using the ArchR addGroupCoverages() function with a minimum of 80 and maximum of 800 cells included per replicate. Peak calling was then performed with MACS2 (v2.2.9.1)^98^ using the addReproduciblePeakSet() function, and a peak-matrix was constructed using addPeakMatrix().

### Integration and Clustering of 10x Genomics Visium HD datasets

Preprocessing, QC, dimensionality reduction, and clustering for spatial transcriptomics datasets were performed using Seurat (v5.1.0). For global clustering, each sample was filtered for spots with >50 unique features. Following QC, log-normalization was performed for each sample using the Seurat NormalizeData() function. Following normalization, the top 2,000 HVGs were identified using the FindVariableFeatures() function. Due to the large number of spatial bins retained for analysis, geometric sketching of the dataset was performed using the Seurat SketchData() function with 50,000 cells and the LeverageScore() method. PCA analysis was performed on the sketched data using RunPCA() with the top 50 PCs retained for downstream integration and clustering. Harmony (v1.2.1)^99^ was used for batch correction on the sketched data prior to nearest neighbor calculations, UMAP dimensionality reduction, and Louvain clustering as described above. Finally, ProjectData() was used to map all retained spatial bins onto the sketched representation to generate the final clustering and UMAP.

### Cell Composition and Single Cell Mapping of Spatial Transcriptomic Datasets Using TACCO

Single-cell clusters were mapped onto spatial transcriptomic datasets with the TACCO package (v0.4.0)^15^ using the tc.tl.annotate() function. All variable names were converted to their respective Ensembl IDs using BioMart annotations to facilitate matching between single cell and spatial transcriptomic datasets. Maximal-likelihood SC annotations were transferred onto the ST dataset using the tc.utils.get_maximum_annotation() function. For single cell-level ST mapping, the same tc.tl.annotate() function was used except with the cell index used as the annotation key. The resulting probability matrix (spot x cell) was used to project cell-level metadata to the corresponding spots in the target spatial transcriptomics dataset.

### Visualization of Spatial Transcriptomic Data Using SpatialData

Spatial transcriptomics datasets were imported using the SpatiaData (v0.4.0)^100^ package with the visium_hd() function to create a Zarr data store. Visualization was performed using the render_image() function to produce the H&E image and render_shapes() function to produce the relevant spatial bins and associated information such as identity or gene expression. For plotting continuous variables, set_zero_in_cmap_to_transparent() was used to set the 0 value in the color map to transparent for visualization clarity. Single mapped populations were visualized by setting the color of all other population identities as transparent (#00000000). Grayscale versions of the H&E image were produced in Adobe Photoshop based on a standard weighted RGB luminance formula.

### Cell Segmentation of Visium HD Spatial Data

Cell segmentation analysis of Visium HD spatial data was performed using bin2cell (v0.3.3)^101^. An anndata object was constructed from the 2µm binned Visium HD spatial data filtering for genes expressed in >3 spots and spots with at least one count. Nuclear segmentation was then performed on the H&E image using the bin2cell stardist() function with a probability threshold of 0.01. Nuclear segmentations were then expanded to neighboring bins using the expand_labels() function. SC-ST mapping of bin2cell segmented cells was performed using TACCO as described above. Visualization of bin-segmented melanocyte populations was performed using the SpatialData package.

### Cell Trajectory and Pseudotime Analysis

Cell trajectory analysis of the matrix and sebaceous gland was performed using Palantir (v1.4.0)^76^. First, the raw and normalized count matrices, dimensionality reductions, and metadata were exported from Seurat and used to initialize a Scanpy anndata object. For the sebaceous gland, spatial bins belonging to the ductal and lobular trajectories were segregated into different objects for calculation of independent trajectories. Diffusion maps for each cell trajectory were obtained using the Palantir run_diffusion_maps() function and the top three components were identified using the determine_multiscale_space() function. Root and terminal nodes were then selected from the component extrema. The run_palantir() function was used to compute pseudotimes using the default parameters. For pseudotemporal analysis of the matrical compartment, two independent trajectories were calculated, one representing the lower COL17A1 compartment and IRS lineage, and another representing the upper COL17A1 compartment and cortex/cuticle lineage. Diffusion maps corresponding to each trajectory and diffusion extrema were calculated as described above. After running Palantir, branch assignments and fate probabilities for each trajectory were calculated using the select_branch_cell() function with default parameters.

### Identification of Gene Expression Trends Across Pseudotime

Gene trends across pseudotime were computed using tradeSeq (v1.22.0)^102^. Expression trends were calculated for genes detected in >5% of bins, with the weights for all bins set to 1. The evaluateK() function was executed using 500 randomly selected genes to determine the optimum number of knots, with 13 knots showing the best performance. A negative binomial generalized additive model (NB-GAM) for gene expression across pseudotime was then calculated using the fitGAM() function. Genes with statistically significant trends across pseudotime were identified using the associationTest() function, with the top 1,500 genes by Wald statistics retained for subsequent analyses. Next, smoothed gene expression values were calculated across pseudotime using the predictSmooth() function. Gene modules were identified through Louvain clustering of the scaled expression values across pseudotime using the FindClusters() function from Seurat. Retained genes were then ordered by point of max expression, and gene trends were visualized using ComplexHeatmap (v2.20.0)^103^. Module GO term enrichment analysis was performed using ClusterProfiler (v4.12.6)^104^. Module scores were computed using AUCell (v1.26.0)^105^ using default parameters.

### Proliferation, Adhesion, and Pigmentation Gene Set Scoring

To analyze trends in cell proliferation, a general proliferation score was computed with AUCell using a previously published S and G2M gene set^106^ embedded in the Seurat package. To calculate the matrix adhesion score, CellChat (v2.1.1)^107^ cell-cell interaction analysis was first performed to identify any relevant receptor-ligand interactions (ECM-receptor category) between the matrix and dermal papilla cells. Cell communication analysis was performed by converting single cell expression matrices into CellChat objects and subsetted for genes involved in signaling interactions. Overexpressed genes and ligand-receptor pairs were then calculated using the identifyOverExpressedGenes() and identifyOverExpressedInteractions() functions. Cell-cell communication networks were then computed using computeCommunProb(), and aggregated with aggregateNet(). Significantly enriched ligand-receptor pairs for each identified cell interaction were extracted using the extractEnrichedLR() function, and genes corresponding to membrane-bound ECM receptors were grouped into a gene set. Adhesion module scores were then computed using AUCell as described above. For adhesion-proliferation comparisons, cells were stratified into high- and low-adhesion groups based on a median split, and differences in proliferation scores were assessed using a two-sided Mann-Whitney U test with Benjamini-Hochberg correction. Differences in proliferation scores across COL17A1 matrical populations were evaluated using a Kruskal-Wallis test, followed by post-hoc pairwise Mann-Whitney U tests with Benjamini-Hochberg correction. Analysis of pigmentation score differences across melanocyte subpopulations was performed using the statistical framework described above. To analyze Ki67 positivity within the ORS subpopulations, we first calculated the total counts of *MKI67* RNA positive and negative cells within each sample. Population-specific proliferation rates were then estimated with a mixed-effects binomial regression model with a logit link using the lme4 package (v1.1.35.5)^108^ with ORS subpopulation modeled as a fixed effect and donor modeled as a random effect. Statistical significance was assessed through Wald tests on pairwise contrasts of the estimated marginal means.

### Live Imaging of Human Scalp Hair

Live imaging experiments were performed as previously described^74^. Data analyses, including single cell tracking, quantification of cell displacement and migratory speed, were performed in Imaris 10.2. Automated cell identification and tracking was used first, and selected cells were manually examined for tracking fidelity.

### Peak-Gene Linkage Analysis and Domains of Regulatory Chromatin (DORC) Computation

Iterative latent semantic indexing (iLSI)-based dimensionality reduction, Harmony batch correction, and nearest neighbor graph computation of the scATAC data were performed prior to metacell computation and peak-gene linkage using ArchR. Metacell aggregation of the multiomics data was performed using SEACells (v0.3.3) ^109^ using the ATAC LSI embedding as input, targeting ∼75 cells/metacell. Gene-peak linkages were computed using the get_gene_peak_correlations() function, using a search space of 100 kb surrounding the gene body. To speed up calculations, genes and peaks detected in fewer than 25 cells were excluded from the analysis. Gene-peak linkages with a nominal p value <0.05 and a correlation coefficient >0.25 were considered significant. For DORC computation, highly regulated genes were selected by plotting the number of peaks per gene against their ranks and estimating a cutoff of >= 8 linked peaks based on the elbow heuristic. The peak-gene linkages for all genes falling above this cutoff were then computed using an expanded 500 kb search space. DORC scores for highly-regulated genes (HRGs) were computed using the approach previously described^75^. First, peak fragment counts were normalized to the fraction of reads falling in peaks for each cell, targeting a sum of 10^6 fragments per cell. DORC scores were then calculated as the aggregated fragment counts across all peaks linked to a gene. Differential DORC score analysis across the upper and lower COL17A1 compartments was performed using a Wilcoxon rank sum test implemented with the Seurat FindMarkers() function. Differential peak analysis was performed for the COL17A1 upper and lower compartments using the ArchR getMarkerFeatures() function with TSS enrichment and log10(fragment count) considered as bias variables. Individual peaks with an FDR-corrected p value <=0.2 and an absolute log2fold change >0.5 were considered differentially accessible.

### chromVAR Motif Deviation Analysis

Motif finding and genome track visualizations were performed using Signac (v1.14.0)^110^. ArchR objects were converted into Signac format using the ArchRtoSignac package (v1.0.5)^111^ prior to downstream analysis. Motif analyses for matrix populations were performed using the JASPAR2020 CORE vertebrate transcription factor motif database^112^. Motif positions across all peaks were computed using the Signac AddMotifs() function. ChromVAR motif deviations were then computed using the Signac RunChromVAR() function and the chromVAR package (1.26.0)^113^. Differential motif activity across the Upper and Lower COL17A1 lineages were identified via Wilcoxon rank sum test using the Seurat FindMarkers() function. Motifs with an absolute mean deviation score difference of >0.5 and a Bonferroni-adjusted p value <0.05 were considered differentially active.

### Analysis of DORC and Gene Dynamics Across Pseudotime

For visualization of DORC-associated genes across pseudotime, log-normalized gene expression, and DORC scores were first min-max normalized over all cells belonging to the cortex/cuticle and IRS cuticle trajectories. Next, normalized expression values were smoothed over each trajectory for visualization of trends using the R stats smooth.spline() function with 3 degrees of freedom. To compute latency, gene expression values and DORC scores were then normalized to each individual trajectory, and chromatin-RNA residuals were computed by subtracting the rescaled DORC scores from the rescaled gene expression values. Spatial mapping of DORC activity and gene expression dynamics through single-cell annotation transfer was performed using TACCO single-cell mapping as described above. For clustering of gene dynamics trends along the cortical trajectory, gene expression and DORC scores were smoothed along cortical pseudotime as described above and normalized to the cortical trajectory.

Identification of basal and terminal program genes was performed using a Wilcoxon rank sum test between the first and last pseudotime deciles. Genes with a Bonferroni-adjusted p value cutoff of 0.01 and an absolute average log2fold change >1.75 were retained for downstream dynamics analysis. Hierarchical clustering of gene dynamics was performed using the Ward.D2 algorithm, implemented in the R stats hclust() function, on a matrix of normalized DORC scores, gene expression, and DORC-RNA residuals and visualized using the ComplexHeatmap package.

### Antibody Staining via Immunohistochemistry and Immunofluorescence

Formalin-fixed paraffin-embedded tissue sectioned at 5um were deparaffinized and rehydrated by heating at 60°C for 1 hour followed by passage through a xylene-ethanol gradient. Heat-induced antigen retrieval and tissue depigmentation was performed in Tris-based antigen unmasking solution (Vector Laboratories) and 0.5% hydrogen peroxide at 80°C for 30 minutes. The sections were then washed and permeabilized in phosphate buffered saline with 0.25% Triton-X for 10 minutes at room temperature followed by blocking with 5% secondary-species specific normal sera in phosphate buffered saline with 0.1% Tween-20 (PBST) for 1 hour at room temperature. Next, the sections were incubated in blocking buffer at 4°C overnight in a humidified chamber with primary antibodies specific to human KRT15 (1:100, Santa Cruz, sc-47697), NPNT (1:100, abcam, ab272549), DIO2 (1:100, Invitrogen, PA5-100474), Ki67 (1:100, Cell Signaling, cat. 9027S), DCT (1:100, Santa Cruz, sc-74439), Melan A (1:100, Santa Cruz, sc-20032), SOX10 (1:100, Cell Marque, cat. 383R), KRT6 (1:100, Biolegend, cat. 606101), Collagen XVII (1:100, abcam, ab184996), GATA3 (1:100, Santa Cruz, sc-268), HOXC13 (1:100, Sigma, HPA051634), and P63 (1:100 dilution, Santa Cruz, sc-25268).

For immunofluorescence, sections were treated after primary antibody incubation with Alexa Fluor 555 Goat anti-Mouse (Invitrogen) and/or Alexa Fluor 647 Goat anti-Rabbit IgG (Invitrogen) diluted at 1:2000 in blocking buffer for 1 hour at room temperature. To visualize cell nuclei, sections were counter-stained with Hoechst and mounted in Fluoromount-G medium (Invitrogen). Imaging was performed using a Nikon A1 confocal fluorescence microscope.

For immunohistochemistry, sections were treated after primary antibody incubation with ImmPRESS HRP Horse anti-Rabbit IgG (Vector Laboratories) and/or ImmPRESS AP Horse anti-Mouse IgG (Vector Laboratories) for 1 hour at room temperature. Signal detection was performed using DAB (Vector Laboratories; SK-4105) and/or Vector Red substrates (Vector Laboratories; SK-5100) for 5-30 minutes at room temperature depending on signal strength. Sections were subsequently washed, counter-stained with hematoxylin, dehydrated through an ethanol-xylene gradient, and mounted in Permount medium (Fisher Scientific).

### Sample Cell Proportion Analysis

Single-cell differential composition analysis of disease compared to control samples was performed using scCODA (v0.1.9)^81^. Stacked barplots visualizing the sample cluster-level composition was performed using viz.stacked_barplot(). Grouped barplots divided by sample source was performed to select the cell population with the lowest variability to use as the reference category (eccrine). Modeling was performed with an FDR of 0.15 due to the relatively small sample size. The log2-fold change of the analyzed clusters was plotted using viz.plot_effects_barplot().

### Immune Cell Identity Annotation

Immune cell cluster annotation was performed using a combination of canonical immune cell marker expression and automated cell type annotation using CellTypist (v1.6.3)^114^. In brief, the Immune_All_Low and Immune_All_High models were imported and used to annotate the immune subclusters using celltypist.annotate() with majority voting. The resulting identities were then further refined using cluster-based differential gene analysis and compared against immune subset markers established in published literature.

### Pseudobulk Differential Gene Expression and Gene Set Enrichment Analysis

For pseudobulk differential gene expression, sample-level expression was obtained using the AggregateExpression() function in Seurat. Differential gene expression was performed using the Seurat implementation of DESeq2 through the FindMarkers() function^115^. The resulting DEG list was filtered for an adjusted p < 0.05 and average log2FC > 0.5. GSEA was then performed with the Fast GSEA package using the fgsea() function for the human hallmark gene set derived from the MSigDB collection^116^.

## Data availability

The accession number for the processed sequencing data reported in this paper is GEO: GSE301777 (scRNA-seq), GEO: GSE301766 (multiome ATAC+GEX), and GEO: GSE301780 (spatial transcriptomics). Raw sequencing data deposited to NIH dbGaP and SRA in compliance with Genomics Data Sharing policies. Additional information and data requests can be made to the corresponding author upon request.

## Code Availability

Custom notebooks and scripts used in the data analysis for this project are available in the GitHub repository (https://github.com/ChristopherStephens21/HHA)

## Acknowledgments

The authors thank participants of the study for their contributions to scientific research. We are grateful to members of the Yi laboratory for comments, discussions, and suggestions, including the contribution of Arpan Das and Dr. Yuheng Fu at the early stage of this project. We also thanks Dr. Stephanie Rangel from the Department of Dermatology Clinical Trial Unit, Dr. Jennifer Wai from the NUSeq Core, and members of the NU SBDRC for technical assistance. The authors acknowledge the support from the Che Family Foundation, NIH R01HD107841, R01AR081103, R01AR071435 and R01AR066703 (to RY), Ghandi Family Foundation, NIH T32 Program in Cutaneous Biology 5T32AR060710, NIH P30AR075049 pilot grant, Dermatology Foundation Dermatologist-Investigator Research Fellowship and Physician-Scientist Career Development Award (EBL), NIH F31 Predoctoral Individual National Research Service Award 1F31AR085379-01 (CMS).

## Author contributions

Conceptualization: EBL and RY

Clinical specimen collection: EBL, MK, AC, JH, VLQ, MLC, RLP, SSY

Experimentation: EBL, JH, DW

Bioinformatics analysis and data presentation: EBL, CMS, RY

Live imaging and analysis: NT, TB, IS, RY

Writing: EBL, CMS, RY

*All authors have read and approved the submitted version of the manuscript

## Competing interests

The authors declare that they have no competing interests.

---

**Extended Data Fig. 1:**
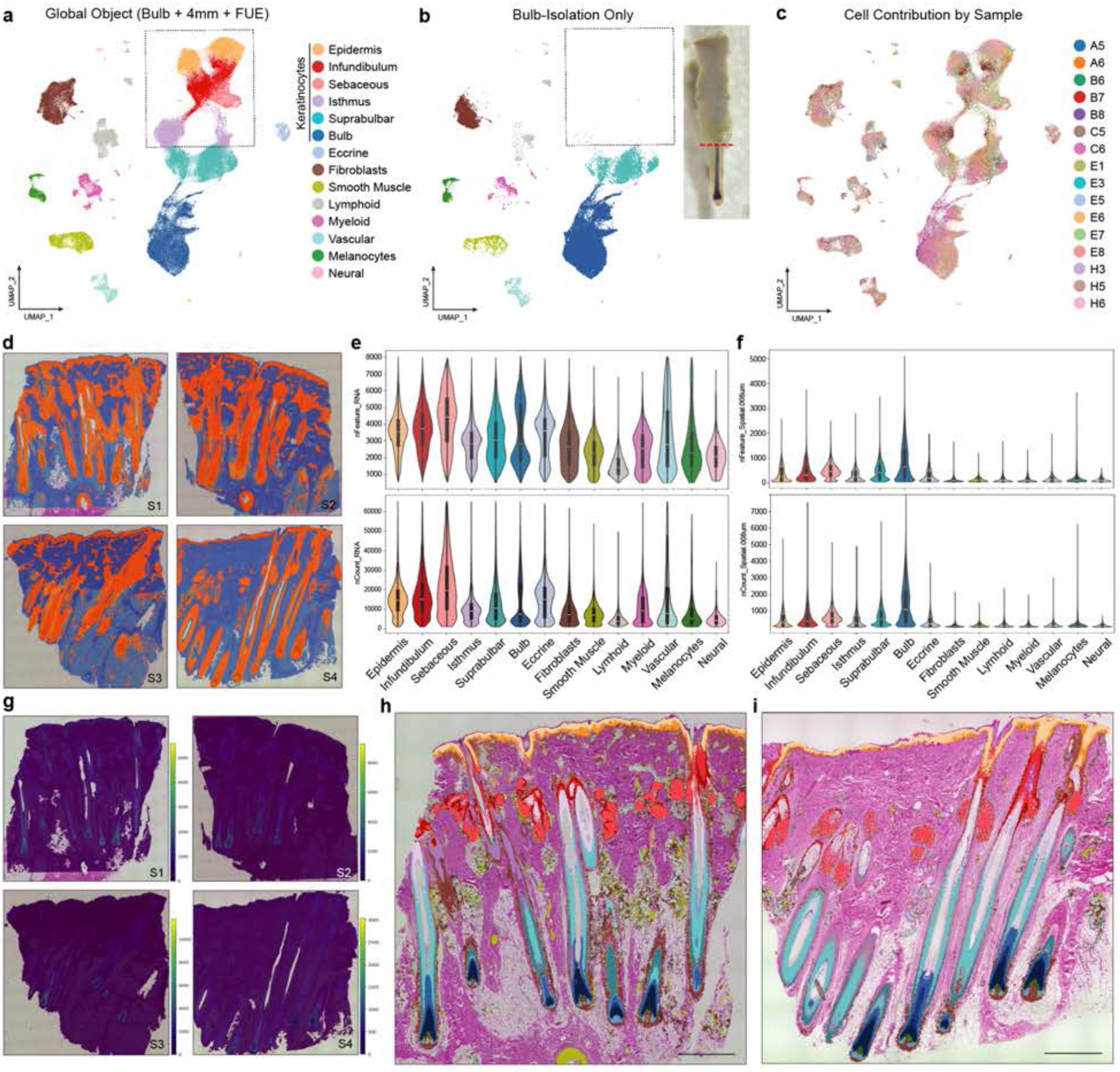
Clustering and Assay Performance Characteristics of Single-Cell and Spatial Transcriptomic Atlas. **(a)** UMAP of global-level clustering for the integrated single-cell RNA-seq and multiomics datasets. **(b)** Single follicular unit microdissection of lower hair follicle/bulb (dashed line) and the cellular contribution of these samples to the global atlas visualized through UMAP. Boxed: no significant contribution of bulb samples to epidermis, infundibulum, sebaceous, and isthmus keratinocytes. **(c)** Cell contributions by sample. **(d)** Visualization of retained (orange) vs. filtered (blue) spatial 8µm bins for ST samples S1-S4. **(e-f)** Violin plots of unique features and UMI counts per cell (e) and per filtered 8µm bins (f), grouped by global atlas-level clusters. **(g)** Spatial heatmap visualization of unique UMI counts per 8µm bins for ST samples S1-S4. **(h-i)** Spatial plotting of atlas-level SC clusters on ST samples S1 (h) and S4 (i). Scale bar: 1mm.

**Extended Data Fig. 2:**
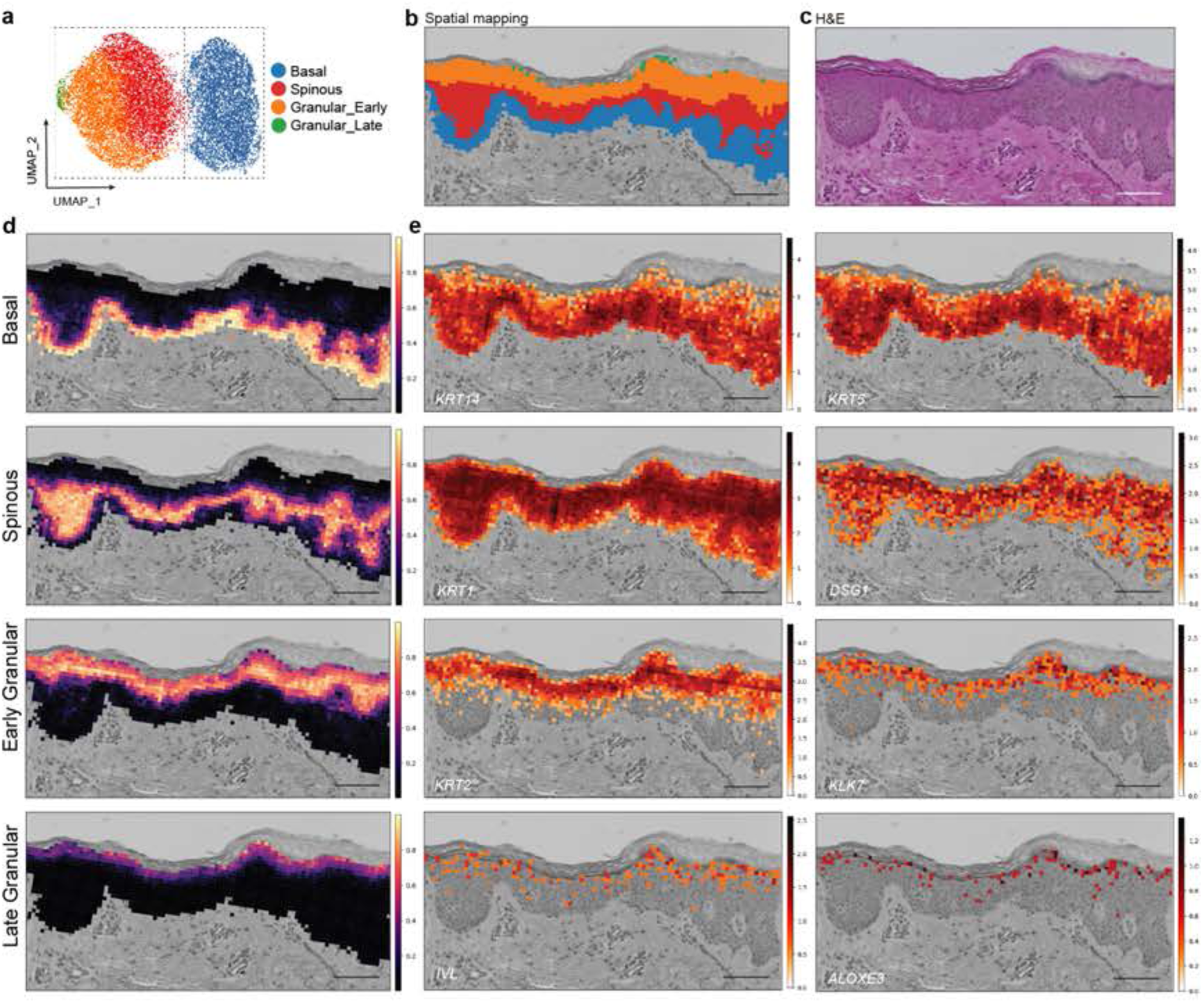
Spatial Mapping of Single-Cell Transcriptome Resolves Canonical Epidermal Architecture. **(a)** UMAP of higher resolution subclustering of IFE cells from SC and **(b)** mapped organization of these IFE subclusters in space showing the canonical layer organization of the epidermis. **(c)** H&E image of the displayed IFE region. **(d)** Spatial heatmap showing probabilistic subcluster spatial distributions rather than categorical bin assignments, with areas of highest probability for each layer being assigned to their respective categorical IFE identities. **(e)** Spatial feature plots of characteristic IFE marker genes by layer. Scale bar: 100µm.

**Extended Data Fig. 3:**
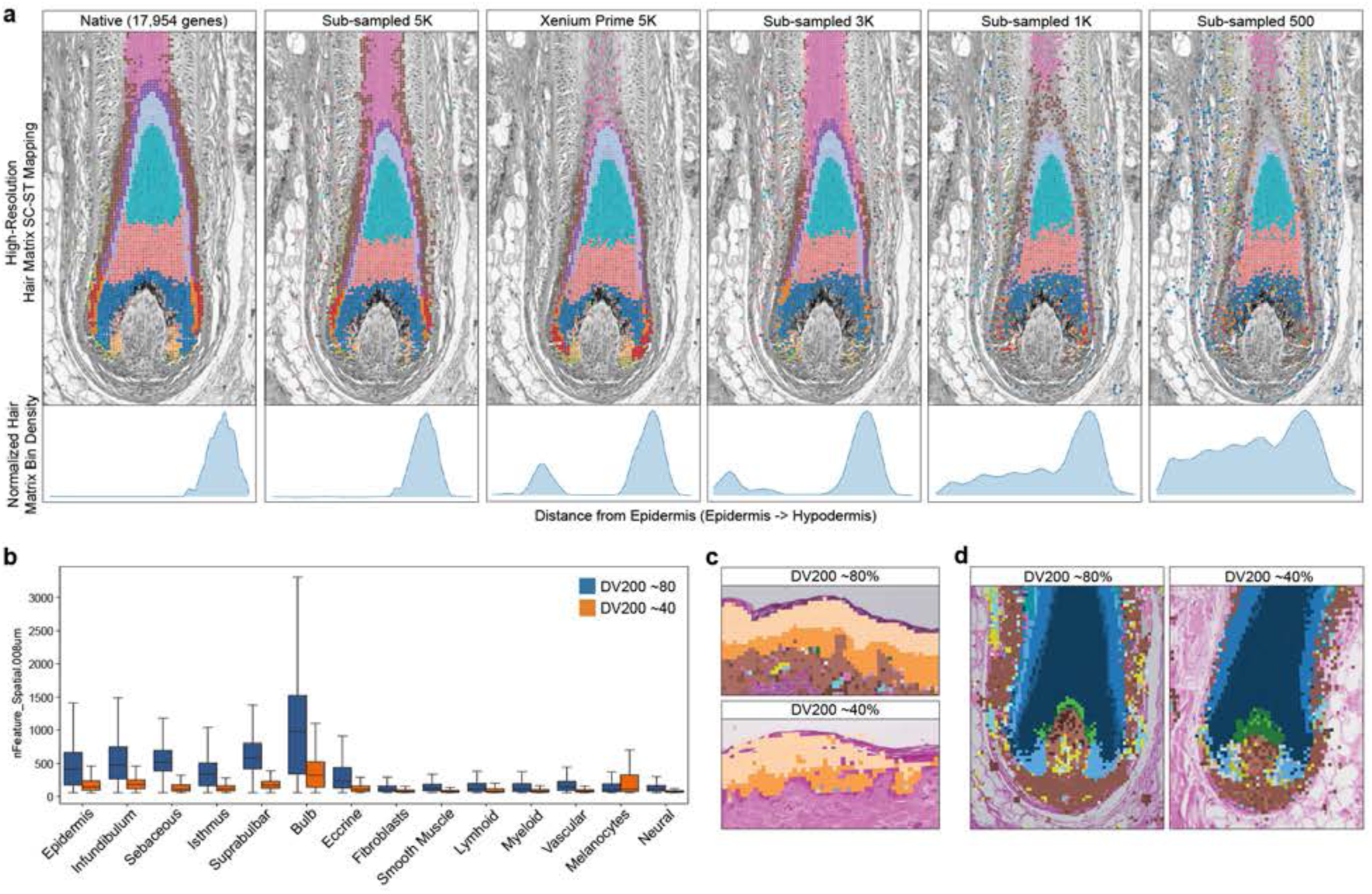
Impact of Gene Capture and Sample RNA Quality on Spatial Mapping Accuracy. **(a)** Comparison of the SC-ST mapping accuracy for high-resolution hair follicle bulb/matrix subpopulations using the native Visium HD, simulated Xenium, and subsampled gene spaces; the bottom panels depict the normalized linear distribution of bins assigned with bulb/matrix identities along the epidermis (left)-hypodermis (right) axis. **(b)** Bar plot comparison of unique genes per 8µm bin captured between DV200 ∼80 and ∼40 samples showing significantly reduced gene detection associated with reduced RNA quality. Cropped image of interfollicular epidermis **(c)** and lower hair follicle/bulb **(d)** showing increased mosaicism with lower RNA quality during SC-ST integration and cluster identity transfer.

**Extended Data Fig. 4:**
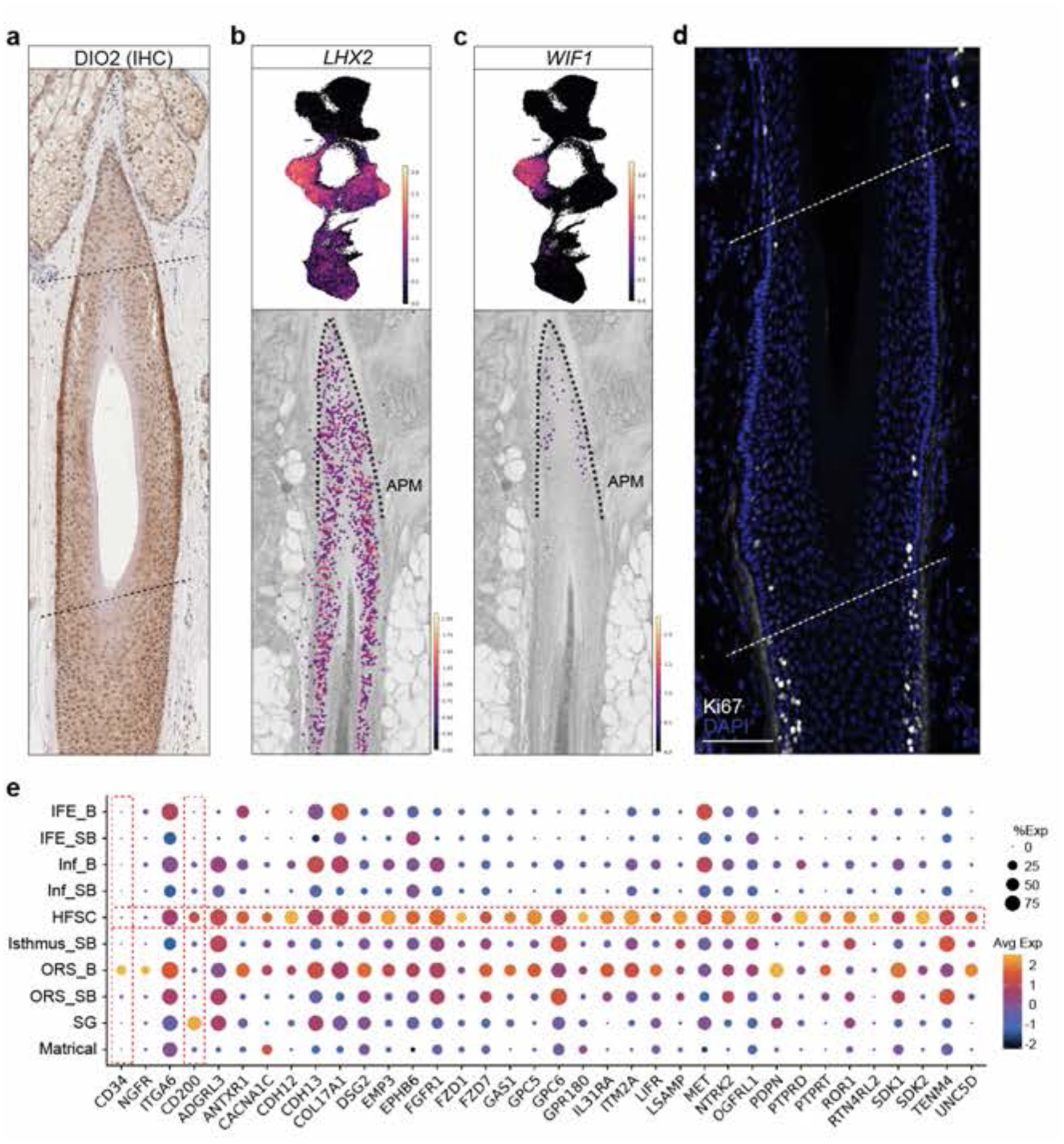
Gene Expression and Cell Proliferation in Human HF-SCs. **(A)** DIO2 immunohistochemistry of the human adult anagen hair follicle showing DIO2-positivity in a spatially restricted HF-SC region below the sebaceous glands. **(b-c)** UMAP feature plots and spatial gene expression visualizations of murine HF-SC markers (*LHX2*) (b) compared to human HF-SC marker (*WIF1*) (c). Dashed line: HF-SC region. APM: arrector pili muscle. **(d)** Ki67 staining of the isthmus and suprabulbar regions of the anagen HF showing the lack of Ki67-positivity in the HF-SC basal layer and decreased positivity in general in the HF-SC region compared to other regions of the ORS. Dashed line: HF-SC region. Scale bar: 100µm. **(e)** Dot plot surface marker gene expression for HF-SC compared to other human scalp skin keratinocyte populations. Boxed genes (*CD34* and *CD200*) represent traditional human HFSC cell sorting markers.

**Extended Data Fig. 5:**
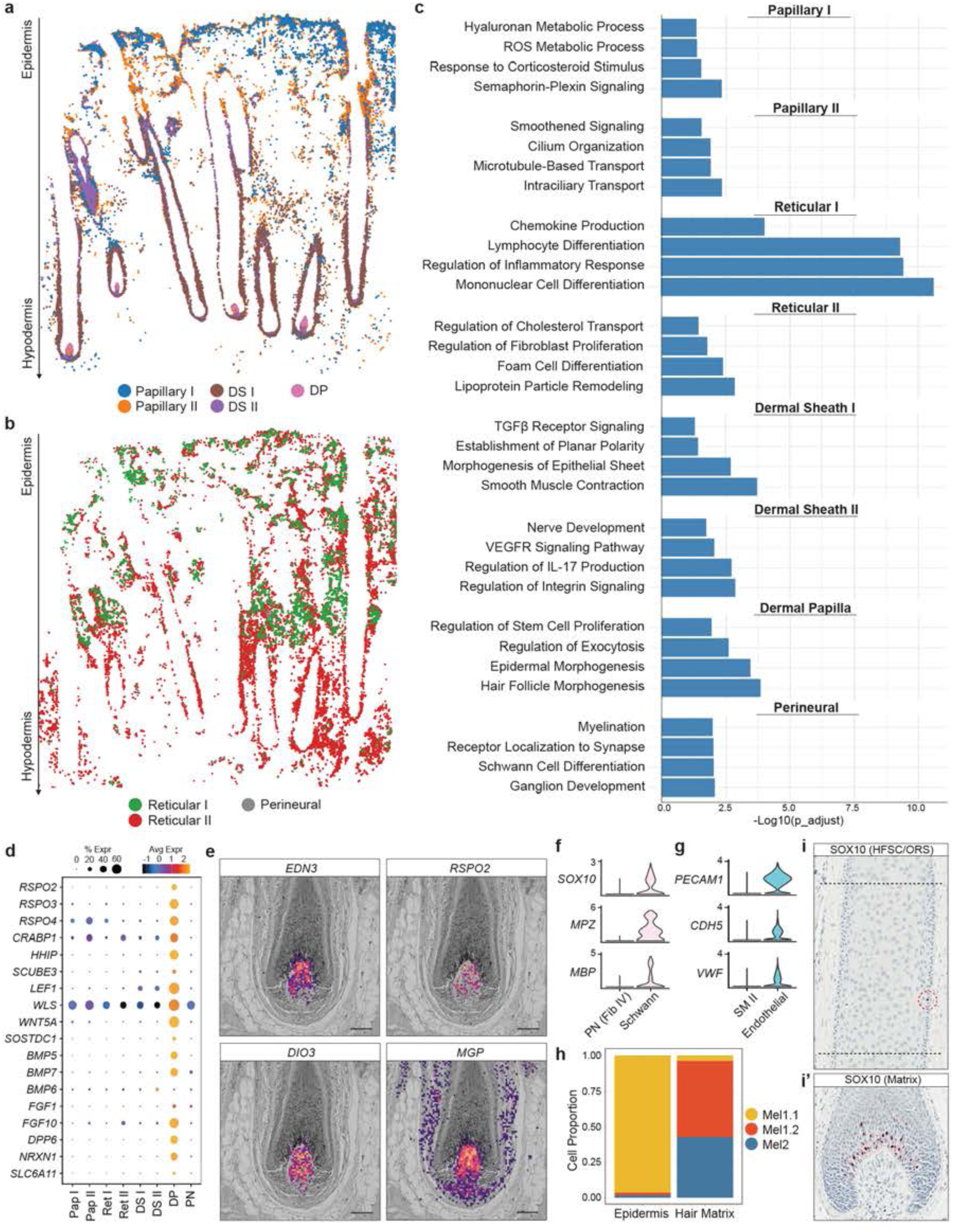
Spatial Mapping Resolves Stromal Cell Heterogeneity. (a-b) Global spatial distribution of fibroblast subpopulations: (a) Papillary and hair follicle associated fibroblast subpopulations and (b) reticular and perineural fibroblast subpopulations. **(c)** Gene Ontology biological process (GO: BP) terms derived from differentially expressed marker genes for fibroblast subpopulation. **(d)** Dot plot of differentially expressed marker genes for the dermal papilla (DP) showing an enrichment of Wnt, BMP, and FGF signaling pathway components. PN: perineural. **(e)** Spatial gene expression feature plots with H&E overlay for select genes (*EDN3, RSPO2, DIO3, MGP*) with distinct intra-DP expression patterns. Scale bar: 100µm. **(f)** Violin plot of canonical Schwann cell markers against perineural (PN) fibroblasts. **(g)** Violin plot of canonical vascular endothelial markers against smooth muscle II (SM II) cells. **(h)** Proportion of spatially mapped cells associated with each melanocyte subpopulation within the IFE and hair bulb showing near-exclusive Mel1.1 distribution in the epidermis and Mel1.2/Mel2 in the hair matrix. **(i-i’)** SOX10 staining of the anagen HF showing rare SOX10-positive cells in the HFSC-region of the ORS (i) and the array of SOX10-positive cells in the upper conical region of the hair matrix abutting the DP (i’).

**Extended Data Fig. 6:**
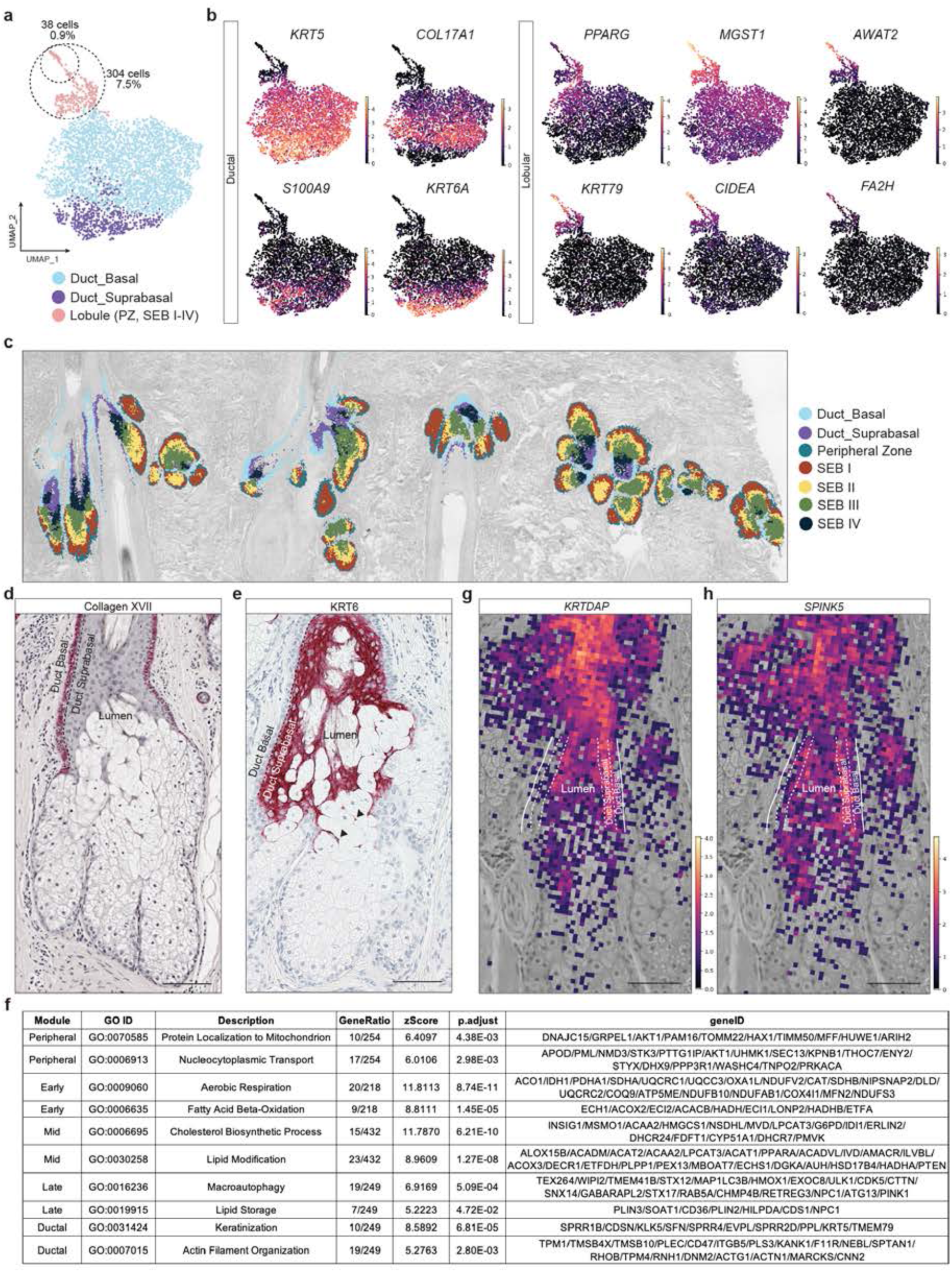
Single Cell Capture of Sebocytes and Spatial Transcriptomic Analysis. **(a)** UMAP of high-resolution subclustering of sebaceous lineage keratinocytes. PZ: peripheral zone. SEB: sebocyte. Inset circles: absolute cell counts and percentage of cells compared to total sebaceous lineage keratinocytes. **(b)** UMAP feature plots for canonical sebaceous gland Ductal (*KRT5, COL17A1, S100A9, KRT6A*) and Lobular (*PPARG, KRT79, MGST1, AWAT2, CIDEA, FA2H*) genes. **(c)** Visualization of high-resolution SG subclusters with H&E overlay across entire ST tissue. **(d-e)** Collagen XVII (D) and KRT6 (E) immunohistochemistry staining of the sebaceous gland shows staining of the ductal basal and suprabasal epithelium respectively. Arrowheads mark filamentous KRT6 positive suprabasal epithelial strands protruding into the SG ductal lumen. **(f)** Details on the featured Gene Ontology terms and their included genes associated with each pseudospatial module for the Lobular trajectory. **(g-h)** Spatial gene expression visualization with H&E overlay for select SG ductal suprabasal genes such as *KRTDAP* (g), and *SPINK5* (h).

**Extended Data Fig. 7:**
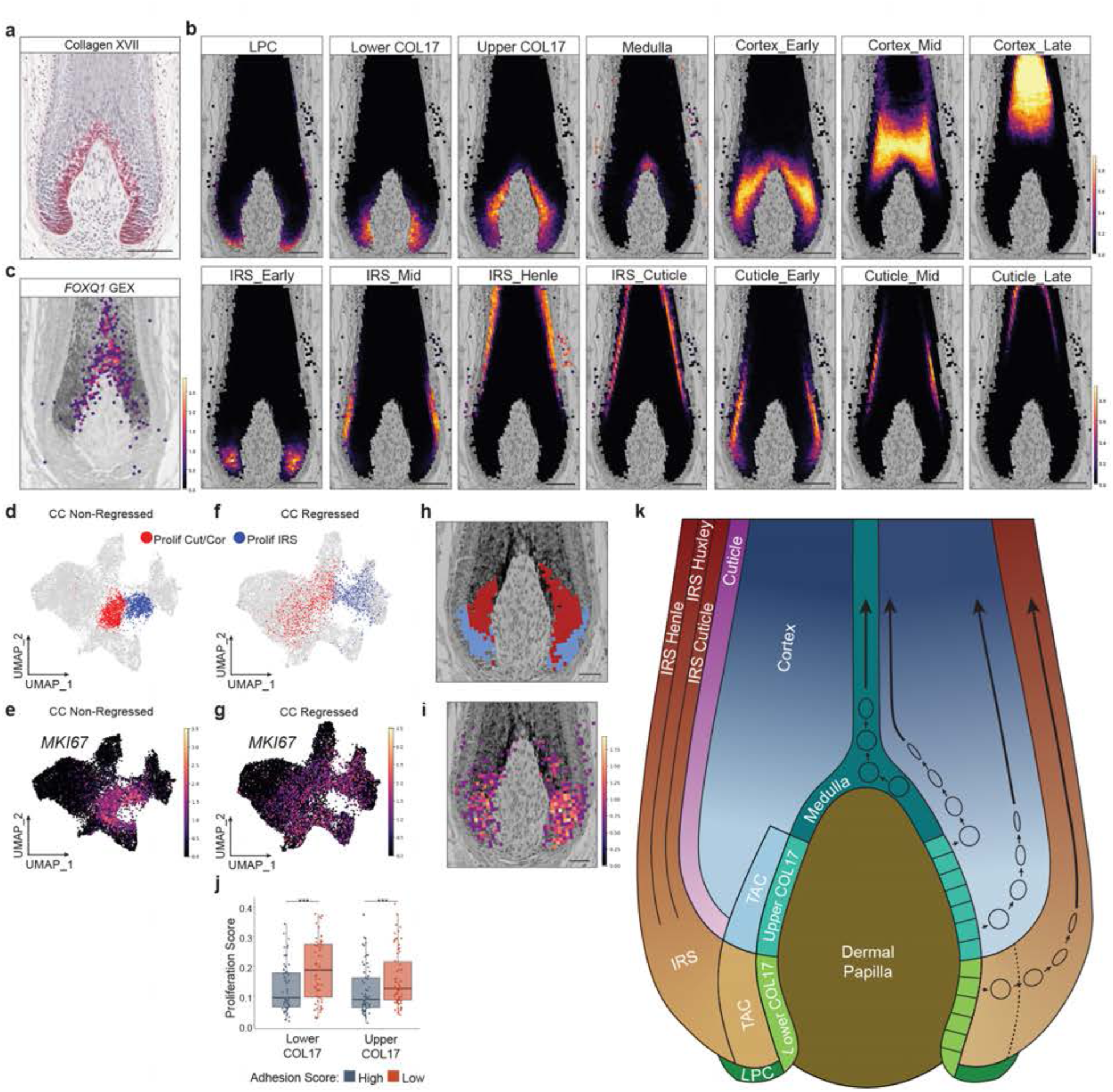
High-Resolution Matrix Population SC-ST Mapping and Cell Cycle Regression on Proliferation Gene Clustering Effects. **(a)** Collagen XVII staining of the HF matrix showing expression restricted to basal cell layers at the matrix-DP junction. **(b)** Spatial heatmap showing probabilistic matrix subcluster spatial distributions rather than categorical bin assignments, with areas of highest probability for each layer being assigned to their respective categorical matrical identities. **(c)** *FOXQ1* spatial gene expression with H&E overlay identifies the medulla lineage. **(d-e)** Centralized UMAP clustering of proliferative matrical cells with no cell cycle (CC) regression (d) and corresponding UMAP feature plot of *MKI67* (e) expression. **(f-g)** UMAP clustering of proliferative matrical cells (f) with cell cycle regression and *MKI67* (g) expression on the regressed UMAP showing the effects of cell cycle regression on the dispersal of proliferative cells to their respective matrical lineages **(h-i)** Spatial mapping of the proliferative matrix cells (blue: proliferative IRS and red: proliferative cortex/cuticle) with H&E overlay (h) and the corresponding matching spatial gene expression pattern of *MKI67* (i) in the hair matrix. Scale bar: 100µm. **(j)** Boxplot of proliferation gene module scores across lower and upper COL17A1 matrical progenitors, divided into low- and high-adhesion cells by their adhesion gene module scores. ***denotes statistical significance from two-tailed Mann-Whitney U test. **(k)** Schematic of hair matrix lineage organization (left) and cellular growth/movement patterns (right).

**Extended Data Fig. 8:**
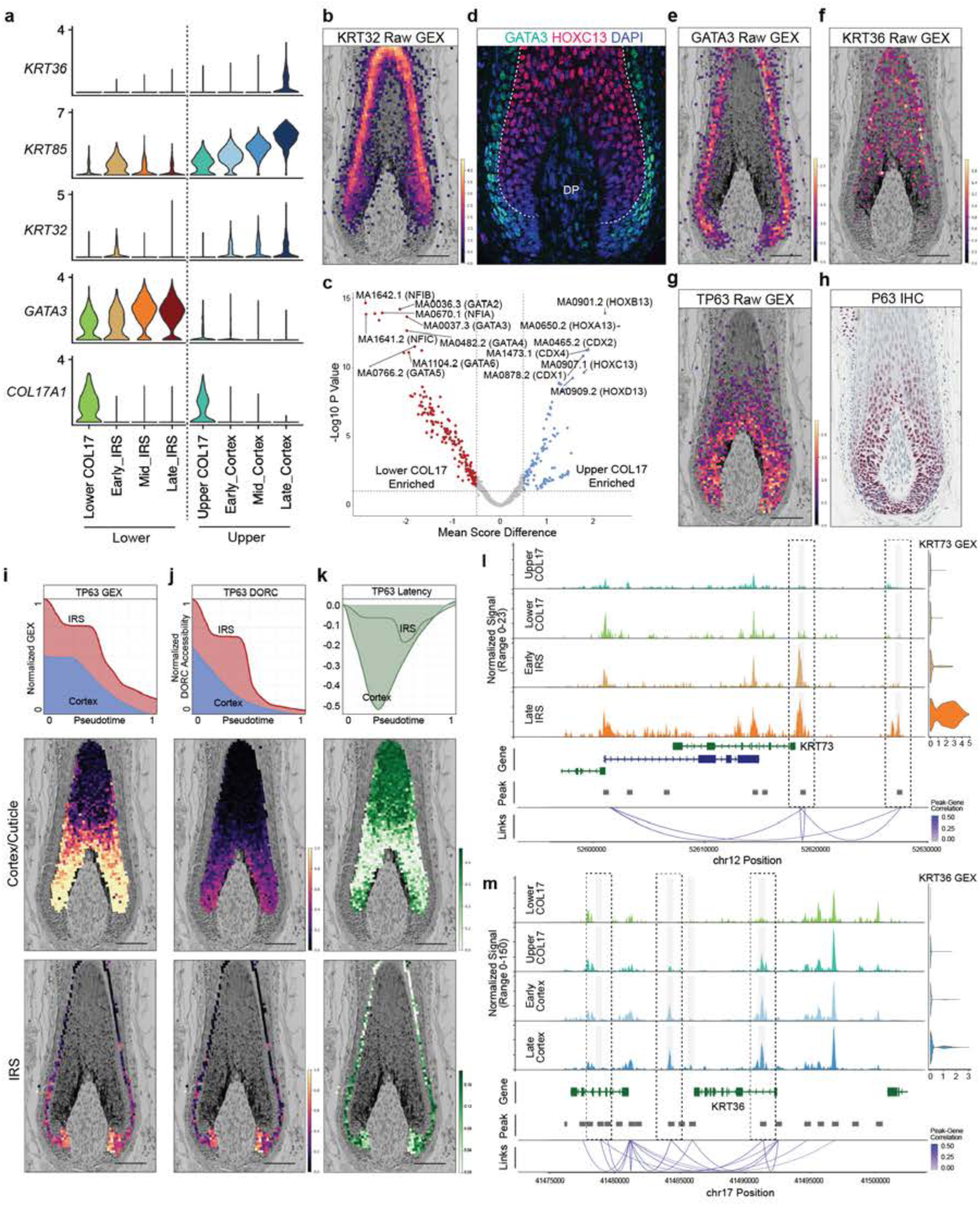
Visualization of Chromatin Dynamics in Gene Expression Priming and Repression. **(a)** Violin plot of COL17A1, lower COL17A1-IRS lineage genes (GATA3), and upper COL17A1-cortex/cuticle genes (KRT32, KRT85, KRT36) expression for the lower and upper lineage cell populations. **(b)** Spatial gene expression visualization of *KRT32.* **(c)** Volcano plot of ChromVAR motif deviation score differences across lower and upper COL17 clusters showing enrichment of GATA and HOXC motifs, respectively. **(d)** GATA3/HOXC13 staining of the hair matrix showing sharp delineation of the lower and upper COL17A1 lineages. **(e-g)** Spatial gene expression visualization of *GATA3* (e), *KRT36* (f), and *TP63* (g) with H&E overlay of the hair matrix. Scale bar: 100µm. **(h)** P63 immunohistochemistry staining of the hair matrix showing progressive downregulation during keratinocyte differentiation. **(i-k)** Line plot of *TP63* gene expression (GEX) (i) and chromatin accessibility (DORC) (j) for the cortex/cuticle and IRS lineages as well as the difference between chromatin accessibly and gene expression (latency) (k). Below each panel is a visualization of the single-cell mapped corresponding values onto ST data with H&E overlay of the hair matrix (middle = cortex/cuticle mapping; bottom = IRS mapping). Scale bar: 100µm. **(l-m)** Peak-to-gene linkage plot of KRT36 (l) and KRT73 (m) showing progressive differential chromatin accessibility peak changes across the lower and upper COL17A1 lineages. Boxed: DORC peaks with significant change with pseudotime. Early and late IRS/cortex defined non-COL17A1 cells below or above the median pseudotime value respectively.

**Extended Data Fig. 9:**
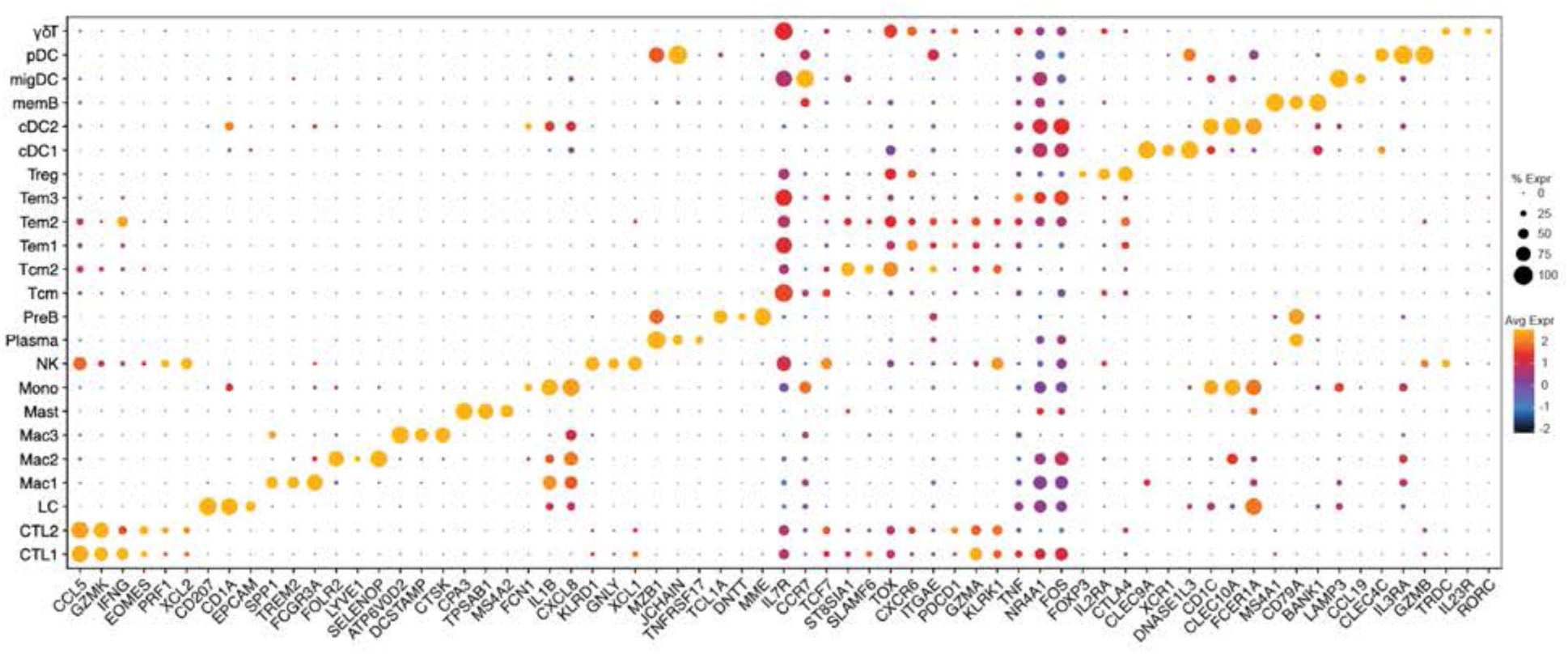
**Immune Cell Cluster Identity and Marker Gene Expression.**

## Notes

https://www.hha.fsm.northwestern.edu

### Competing Interest Statement

The authors have declared no competing interest.

